# Growth dynamics and genetic variation in attached and free-living populations of the filamentous brown seaweed, *Pilayella littoralis*

**DOI:** 10.1101/2023.05.31.543056

**Authors:** Steven L. Miller, Robert T. Wilce

## Abstract

We used common garden growth experiments to study genetic variation among geographic isolates (Greenland, Massachusetts, and Connecticut, USA) of the filamentous brown seaweed *Pilayella littoralis,* including the free-living form unique to Nahant Bay, Massachusetts. Ecotypic variation for temperature growth maximum was demonstrated for a west Greenland isolate (10° C *versus* 15° C for other attached isolates) and between Nahant Bay attached (narrower phenotypic plasticity) and free-living forms (broader phenotypic plasticity) of the species. Morphological and reproductive characteristics of attached and free-living isolates remained distinctive under identical culture conditions after four years. The attached forms were characteristically cabled, twisted, and clumped; unilocular reproductive cells were common and plurilocular reproductive cells were present. The free-living form was characteristically loosely branched and ball-like; only vegetative reproduction occurred, with a few unilocular reproductive cells observed in one experiment. Free-living and attached isolates cultured using no water movement and turbulent conditions to mimic surf and surge conditions did not develop forms that resembled each other after eight months. We additionally used starch gel electrophoresis to study genetic variability in attached and free-living forms of *P. littoralis* from Nahant Bay. Free-living and attached populations were not different at the isozyme level because a limited number of isozymes were resolved (six out of 39 enzymes tested). One isozyme (PGI) was polymorphic, with two alleles present. The two alleles shared in the attached and free-living populations suggest that the free-living form is not one large identical clone. For attached and free-living *P. littoralis,* both transplant and growth studies in the laboratory provide convincing evidence of ecotypic differentiation.

## Introduction

*Pilayella littoralis* (L.) Kjellman is a common brown alga in the North Atlantic. The species ranges south from the high Arctic to New Jersey and Portugal. The alga occurs in brackish waters at the head of estuaries and in fully marine habitats (Russell, 1963; Bolton, 1979) where it attaches to rocks and other algae throughout the littoral and upper sublittoral zones. Individuals of the species are typically erect, uniseriate, with opposite branching usually evident on central axes that exhibit basal-distal polarity. Many studies document the variable morphology and phenology of the species (Kjellman, 1890; Kuckuck, 1891; Holmes, 1893; Knight, 1923; Taylor, 1957; Russell, 1963; Wilce et al., 1982; Pedersen, 1984), including a distinct free-living form in Nahant Bay, Massachusetts (Wilce et al., 1982, 1983; Quinlan et al., 1983, Miller 1988). Characteristics that distinguish the free-living form include radial branching resulting in a ball-like morphology, reproduction by fragmentation of filaments rather than by gametes or spores, polystichous development in central axes, and population persistence year-round.

The wide geographical distribution of *Pilayella littoralis,* its euryhaline characteristics, and variable morphology suggests that 1) the alga exhibits sufficient phenotypic plasticity (*sensu* Bradshaw, 1965) to adapt to environmental conditions throughout its geographic range, or 2) populations are characterized by distinctive phenotypes that persist based on genetic polymorphisms found within the species. Turesson (1922) coined the term ecotype to describe genetically different strains of a species restricted to specific habitats. The classic work of Claussen et al. (1948) on the perennial herb *Achillea millefoliun* (L.) illustrates ecotypic differentiation particularly well.

Determining factors affecting the geographic distribution of marine algae is an area of active research. Growth and reproductive responses to effects of light, temperature, and photoperiod in laboratory studies often correlate with the known geographic distribution of seaweeds (Edwards 1970, 1979; van den Hoek, 1975, 1982, 1984; Lawson, 1978; Kapraun, 1980; Santelices, 1980; Yarish et al., 1984; South, 1987), but few have investigated the temperature and photoperiod responses of isolates of seaweeds collected throughout their latitudinal distribution (Bolton and Luning, 1982; Bolton, 1983; McLachlan and Bird, 1984; Rietema and van den Hoek, 1984). Broad phenotypic variation is common (reviewed by Norton et al., 1981) and can correlate with temperature (Luning et al., 1978; Chen and Taylor, 1980; Bolton, 1983; Novaczek, 1984), salinity (West, 1972; Russell and Bolton, 1975; Bolton, 1979; Reed and Russell, 1979; Yarish et al., 1979; Yarish and Edwards, 1982; Russell, 1985), and other chemical factors. Additional environmental parameters that correlate with ecotypic differentiation in algae include heavy metals (Russell and Morris, 1970; Stokes et al., 1973; Hall, 1980; Shehata and Whitton, 1982; Reed and Moffatt, 1983), nutrients (Francke and Cate, 1980; Espinoza and Chapman, 1983), photoperiod (West, 1972: Luning, 1980a,b), and wave action (Paula and Oliveira, 1982).

Some studies failed to find ecotypic differentiation attributable to temperature among populations of algae despite large differences in temperature between collection sites (Bolton and Luning, 1982; McLachlan and Bird, 1984; Rietema and van den Hoek, 1984; Stewert, 1984). These studies are interesting because water temperature is the primary parameter that correlates with species distributions in the marine environment (Setchell, 1915, 1920; Edwards, 1970, 1979; Edwards and Kapraun, 1973; van den Hoek, 1975; 1982; Kapraun, 1978).

We investigated phenotypic plasticity and genetic differentiation among geographical isolates of *Pilayella littoralis* from Greenland to Connecticut, including the Nahant Bay free-living form. We used cross-gradient culture studies and common garden transplants to evaluate how light and temperature affect the growth, morphology, and reproduction of attached and free-living forms of the species. However, determining genetic variation by measuring growth responses to varied environmental parameters or transplant experiments depends on the often-difficult task of measuring phenotypic responses. To get around these problems, we used starch gel electrophoresis to assess isozyme variability (Lewontin and Hubby, 1966; Clegg and Allard, 1972; Powell, 1975; Gottlieb, 1977, 1981; Nevo, 1978, Hamrick and Holden, 1979; Hamrick et al., 1979; Silander, 1979; Burton, 1983). Genotypes can be identified by electrophoresis based on allele distributions at polymorphic loci, rather than by morphological appearance, which may reflect phenotypic plasticity rather than genetic diversity. Among seaweeds, electrophoretic investigations of genetic variability are uncommon (Malinowski, 1974; Cheney and Babbel, 1978; Miura et al., 1979; Innes and Yarish, 1984; Innes, 1987).

## Methods

### Cross-gradient culture studies

We studied the growth responses of attached and free-living *Pilayella littoralis* to various irradiance, temperature, and photoperiod combinations using a cross-gradient (temperature x light) culture apparatus (modified after Siver, 1983). Three distinct Nahant Bay populations of *P. littoralis* were investigated, including 1) an epiphytic annual population from Galloupes Point, 2) an epilithic perennial population from Dread Ledge, and 3) a free-living perennial population found throughout the bay (Figure 1). Two additional populations studied included 4) an epilithic population from Disko Island, west Greenland, and 5) an epiphytic (on *Ascophyllum nodosum* (L.) Le Jolis) population from a warm water site in southern Connecticut (Waterbury, near the Millstone nuclear power plant). Collection sites, dates of collections, and isolate numbers are in Table 1.

**Figure 1.**
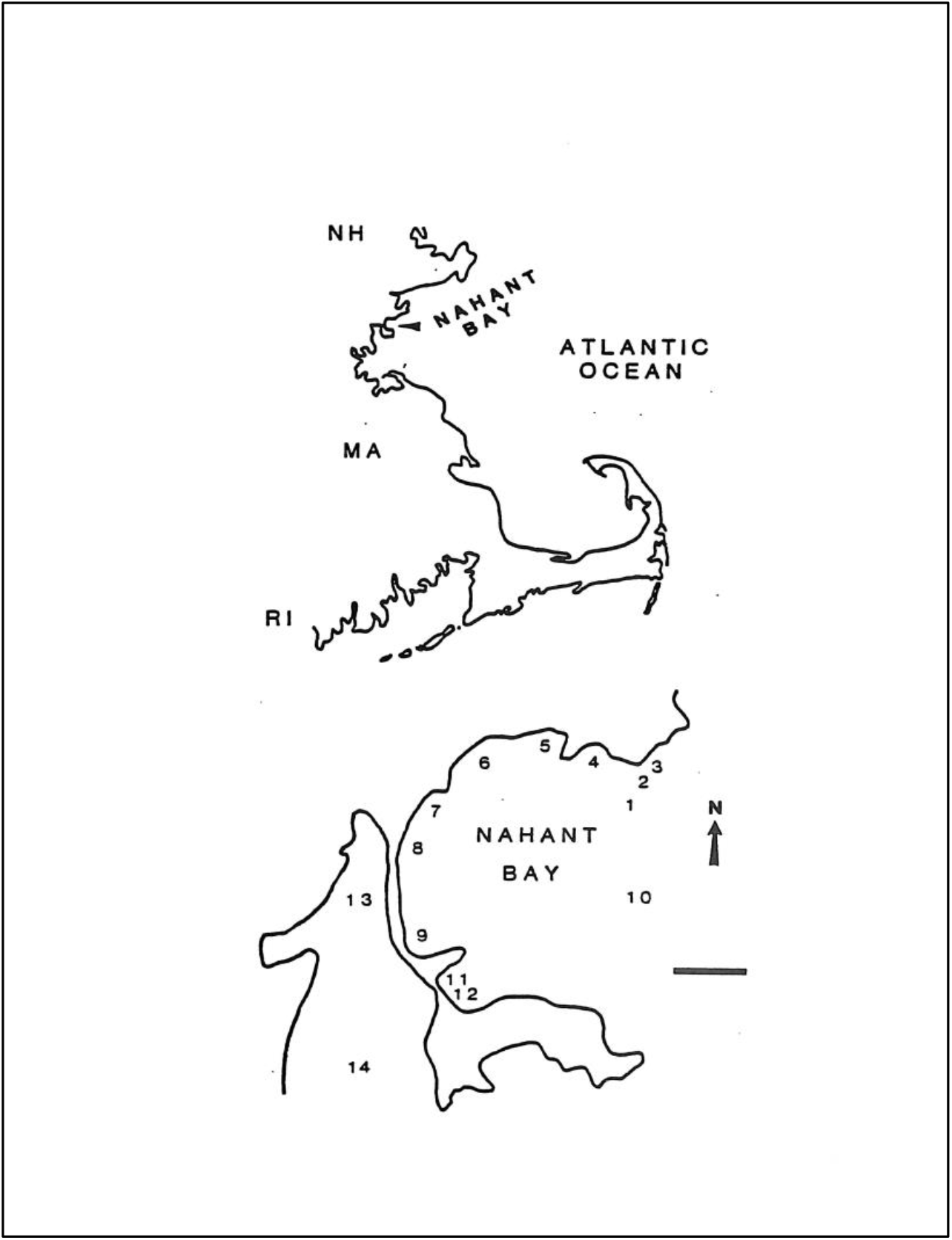
Map of Massachusetts coastline and study sites at Nahant Bay. Numbered sites: 1) Swampscott sewage outfall; 2) Dread Ledge; 3) Galloupes Point; 4) Whale’s Beach; 5) Fisherman’s Beach; 6) King’s Beach; 7) Long Beach - 300 Lynn Shore Drive; 8) Long Beach - MDC House; 9) Long Beach - Tides Restaurant; 10) Nahant Bay Center; 11) Little Nahant North; 12) Little Nahant Coast Guard Beach; 13) Lynn Harbor; 14) Broad Sound - Revere. Bar = 1 Km.

**Table 1.** Sites, dates of collection, and isolate numbers for *Pilayella littoralis* used in cross-gradient growth studies.

| Isolate Identification | Collection Date | Site Location |
| --- | --- | --- |
| Free-living<br>I-143 | February 1984 | Nahant Bay, MA |
| Free-living<br>III25 | June 1984 | Nahant Bay, MA |
| Attached<br>III-9<br>III-75<br>III-3 | June 1984<br>June 1984<br>July 1984 | Nahant Bay, MA<br>Galloupes Point |
| Attached<br>DL-98 | July 1985 | Nahant Bay, MA<br>Dread Ledge |
| Attached<br>GR-29 | October 1986 | Greenland<br>Disko Island |
| Attached<br>CN-13 | June 1987 | Connecticut, USA<br>Waterbury |

To minimize acclimation before collection, we cultured algae under identical temperature (12.5° ±2° C), irradiance (10 uE/m^2^/s), and photoperiod (long days; 16:8, or short days 8:16) for between two months and three years before conducting experiments. Apical cuttings approximately 1 mm long were taken from Nahant Bay free-living and attached plants cultured at least ten months before the start of each experiment. In addition, material from Greenland and Connecticut was cultured for 2-12 months before initiating investigations. Clonal cultures, derived from apical cuttings, were grown to provide inoculum for use in the experiments.

Ten apical cuttings, approximately 1 mm long, were used to inoculate each culture. The fastest-growing five tips in each treatment were retained at the end of one week since not all apical cuttings exhibited growth. Algae were cultured in Pyrex storage dishes containing 250 ml of enriched seawater medium (F/2 or F/4; modified after Guillard and Ryther, 1962). The culture medium in all vessels was changed every 3-7 days. Light was from two General Electric F72T12-CW high-output fluorescent tubes. A Li-Cor quantum meter measured photosynthetically active radiation, 400 - 700 nm. Three irradiances were used in each cross-gradient study: 60±5, 30±5, and 15±5 uE/m^2^/s. Variability in irradiance was due to rotating positions of culture vessels in paired experiments. Algae were grown under two photoperiods, long days at 16 hours light and 8 hours dark, and short days at 8 hours light and 16 hours dark. Up to seven temperature treatments were used on the cross-gradient apparatus, ranging from 0° C to 30° C in 5-degree increments. The temperature did not vary within any treatment by more than ± 1° C. Algae that did not appear to survive a treatment were transferred to 12.5° ±2° C for up to one month to check for viability.

For *Pilayella littoralis* collected from Nahant Bay, an attached isolate and a free-living isolate were usually run in pairs on the cross-gradient apparatus to facilitate comparisons between the two forms of the species. This minimized variation among individual cross-gradient studies. Following each cross-gradient culture study, algae were photographed, microscopic observations were made of reproductive condition and general morphology, and dry weights (to constant weight at 60° C) were determined for each treatment. Growth data were used to test the null hypothesis of no difference in growth response among *P. littoralis* collected from several locations throughout its geographical distribution.

Based on a growth study described below, free-living *Pilayella littoralis* exhibited exponential growth under conditions similar to cross-gradient cultures. Therefore, specific growth rates were calculated as percent increase per day after Guillard (1973) and Brinkhuis (1985). Specific growth rate data were converted to biomass doubling rates (in terms of days). Specific growth rates were also calculated for attached isolates of *P. littoralis,* but confirming exponential growth was problematic because not all growth was vegetative (reproductive organs and germlings were commonly produced). Because dry weights of individual apical cuttings are challenging to measure accurately, inoculum weight was estimated by taking the weight of 250 apical cuttings approximately 1 mm long and replicated three times. The weight of one apical cutting averaged 0.6 ug dry weight.

Ten apical cuttings (approximately 1 mm long) were added to each of 18 glass storage dishes, and growth was measured after one month. Culture conditions were 10° ±1°C, long days (18:6; light:dark), and 30±5 uE/m^2^/s. At each sample interval, algae from three dishes were rinsed in distilled water to remove salts and then dried to constant weight at 60° C and weighed.

### Effects of environment on morphology

To distinguish between environmental and genetic components affecting the morphology and reproduction of *Pilayella littoralis,* attempts were made to modify growth forms by manipulating their environments. Clones derived from attached *P. littoralis* were grown in bubbled cultures to produce morphologies resembling the free-living alga. Actively growing cultures (see below) were used to inoculate 4-liter aspirator bottles with a continuous stream of bubbles entering from the bottom or without bubbles (static conditions). In bubble culture, algae were tossed about in turbulent conditions to mimic surf and surge. Conversely, free-living *P. littoralis* was cultured under bubbled and static conditions. Cultures were started with apical tips (approximately 1 mm long), and subcultures were made over an eight-month period. As bubble cultures of attached clonal material developed, algae with morphologies that resembled the free-living type were selected to start subcultures. These included 1) detached germlings that developed from motile cells produced by the original inoculum, 2) small rounded unattached tufts with a dense central region that developed from precocious germination of sporangia, and 3) pieces of algae that had a distinct axial region.

Apical cuttings were also taken from the above three growth forms to inoculate additional bubble cultures. On one occasion, a bubble culture was started with motile cells derived from unilocular sporangia from an attached plant. These motile cells were carefully collected by pipet with an inverted microscope to ensure no vegetative fragments were included in the inoculum. Static cultures and subcultures of material derived from free-living *P. littoralis* were terminated after two months. Algae were collected and preserved in 5% formalin throughout these studies to assess morphology and reproduction. Photographic records were made. Statistical tests included ANOVA, Scheffe’s Multiple Range Test, and comparing means and their 95 percent confidence intervals.

### Electrophoresis

Horizontal starch gel electrophoresis techniques were modified after May et al., (1979), Shields et al., (1983), Cheney (1985), and Werth (1985). Plexiglass strips (1.2 cm thick) were clamped to glass plates to provide forms (12.5 cm x 21.5 cm) into which starch gels were poured. Gels were prepared using a mixture of 32 g hydrolyzed potato starch (Electrostarch Company, Madison, Wisconsin) and 3 g Connaught’s hydrolyzed starch (Fisher #S676-2), dissolved in 250 ml buffer (Table 2). The gel suspension was prepared as in May et al., (1979), poured into one gel form, and cooled overnight at room temperature.

**Table 2.** Electrophoresis gel and electrode buffer systems.

| Buffer Type | Electrode Buffer | Gel |
| --- | --- | --- |
| Lithium hydroxide<br>(Ridgeway et al. 1974) | LiOH 0.03 M<br>Boric acid 0.30 M<br>pH = 8.1 | 1 ml of electrode buffer to<br>100 ml of TRIS-citrate<br>buffer<br>TRIS 0.03 M<br>Citric acid 0.005 M |
| TRIS-borate-EDTA<br>(Ayala et al. 1974) | Gel and Electrode Buffers<br>TRIS 0.087 M<br>Boric acid 0.0087 M<br>Na<sub>2</sub>-EDTA 0.001 M<br>pH = 9.1 |  |
| TRIS-borate-EDTA<br>(Markert and Faulhaber<br>1965) | Stock:<br>TRIS-base 0.9 M<br>Boric acid 0.5 M<br>Na<sub>2</sub>-EDTA 0.02 M<br>pH = 8.5 |  |
|  | Dilutions:<br>Anodal 1:7<br>Cathodal 1:5 | 1:20 dilution of electrode<br>buffer |
| Amine citrate<br>(Clayton and Tretial 1972) | Citric acid 0.04 M<br>Adjust pH = 6.1 with N-(3-aminopropyl)-morpholine | 1:10 dilution of electrode<br>buffer |
| TRIS-citrate (Soltis et al<br>1983) | TRIS 0.223 M<br>Citric acid 0.069 M<br>pH = 7.2 | Bring 35 ml of electrode<br>buffer to 1 liter with D-H<sub>2</sub>O<br>pH = 7.2 |
| Discontinuou TRIS-Citrate<br>(Poulik system in Soltis et al<br>1983) | NaOH 0.1 M<br>Boric acid 0.3 M<br>pH = 8.6 | TRIS 0.015 M<br>Citric acid 0.004 M<br>pH = 7.8 |

Enzyme extracts were prepared by grinding approximately 0.25 g (blotted wet weight) of algae to a fine powder in liquid nitrogen. The frozen powder was immediately suspended in approximately 0.25 ml of extraction buffer (Table 3) modified after Marsden et al., (1981). Debris was sometimes removed from the homogenate by centrifugation, but usually samples were prepared by directly applying the homogenate to one end of a small rectangular filter paper wick (Whatman #1). Centrifugation did not improve the smearing of lanes that occurred in some enzyme systems. Up to 40 wicks were loaded into a gel in a slit cut lengthwise 3 cm from the cathodal end. The wicks were removed after 15 minutes of electrophoresis at high voltage (to load enzyme extract into the gel), and the two sections of gel were then carefully and firmly placed back together. Electrophoresis was carried out at 4° C, with an ice pack on top of the gel to help maintain cold, even gel temperatures. After completion of electrophoresis, determined by the movement of a dye marker (red food coloring), the gel was sliced 4-5 times horizontally using a series of 1.6 mm plastic spacers to guide nylon monofilament thread through the gel. Each gel slice was individually placed into a staining tray. Staining followed the recipes of Brewer (1970), Shaw and Prasad (1970), Harris and Hopkinson (1976), and Soltis et al. (1983).

**Table 3.** Extraction buffer (Modified after Marsden et al. 1981, 1984) used for electrophoresis of *Pilayella littoralis*. Prehydrate PVPP overnight in buffer; MW = Molecular Weight; Adjust pH = 7.2-7.4.

| Chemical | MW | mM | g/l |
| --- | --- | --- | --- |
| MOPS | 209.3 | 50 | 10.46 |
| L-Ascorbic acid | 336.2 | 2 | 0.62 |
| CaCl<sub>2</sub> | 147.0 | 5 | 9.9 |
| TWEEN 80 |  |  | < 1 ml/l |
| MgCl<sub>2</sub> | 203.3 | 5 | 1.02 |
| DTT | 154.4 | 10 | 1.54 |
| BME | 78.1 | 10 | 0.78 |
| PVP-40 | 10,000 | 2 | 20.0 |
| PVPP |  |  | 100.0 |

Algae from cultured clones were used for electrophoretic analysis. Collections were made at Long Beach of free-living *Pilayella littoralis* and from Galloupes Point of attached *P. littoralis* (Figure 1). Both collections were made in July 1984. Free-living balls were collected individually from the surf zone and were kept in separate containers to prevent the mixing of putative genotypes if the balls fragmented. Attached algae were collected from tufts of material growing on *Ascophyllum nodosum;* one tuft was collected from an *A. nodosum* at intervals of approximately one meter along the shore. Following collection, unialgal cultures were established by repeatedly growing material from apical cuttings (approximately 1 mm long) until all nonbacterial contaminants were removed. Thirty-nine enzymes were assayed, and up to 40 individuals were tested per enzyme. The null hypothesis was no difference in isozyme variation between populations. Alternative hypotheses were that the attached and free-living populations are genetically distinct and that the free-living population is one large clone.

## Results

### Cross-gradient growth studies

To facilitate comparisons of optimal growth temperature among isolates, growth at each temperature is represented as a proportion of total growth under high irradiance. Interactive effects between light and temperature occurred in only one instance (see below). In general, isolates of *Pilayella littoralis* from Greenland, Massachusetts, and Connecticut exhibit growth maxima that correlate with maximum seawater temperature at their respective sites (Figure 2). Specifically, the isolates of attached *P. littoralis* from Nahant Bay exhibited maximum growth at 15° C (Figure 3). For free-living *P. littoralis,* a growth peak was not discernable (with one exception, see below), as considerable growth occurred from 10 - 20° C (Figure 4). In four paired cross-gradient growth studies (utilizing four isolates of attached algae and two isolates of free-living material) growth responses of attached and free-living forms of the species exhibited statistically significant differences (Figure 5).

**Figure 2.**
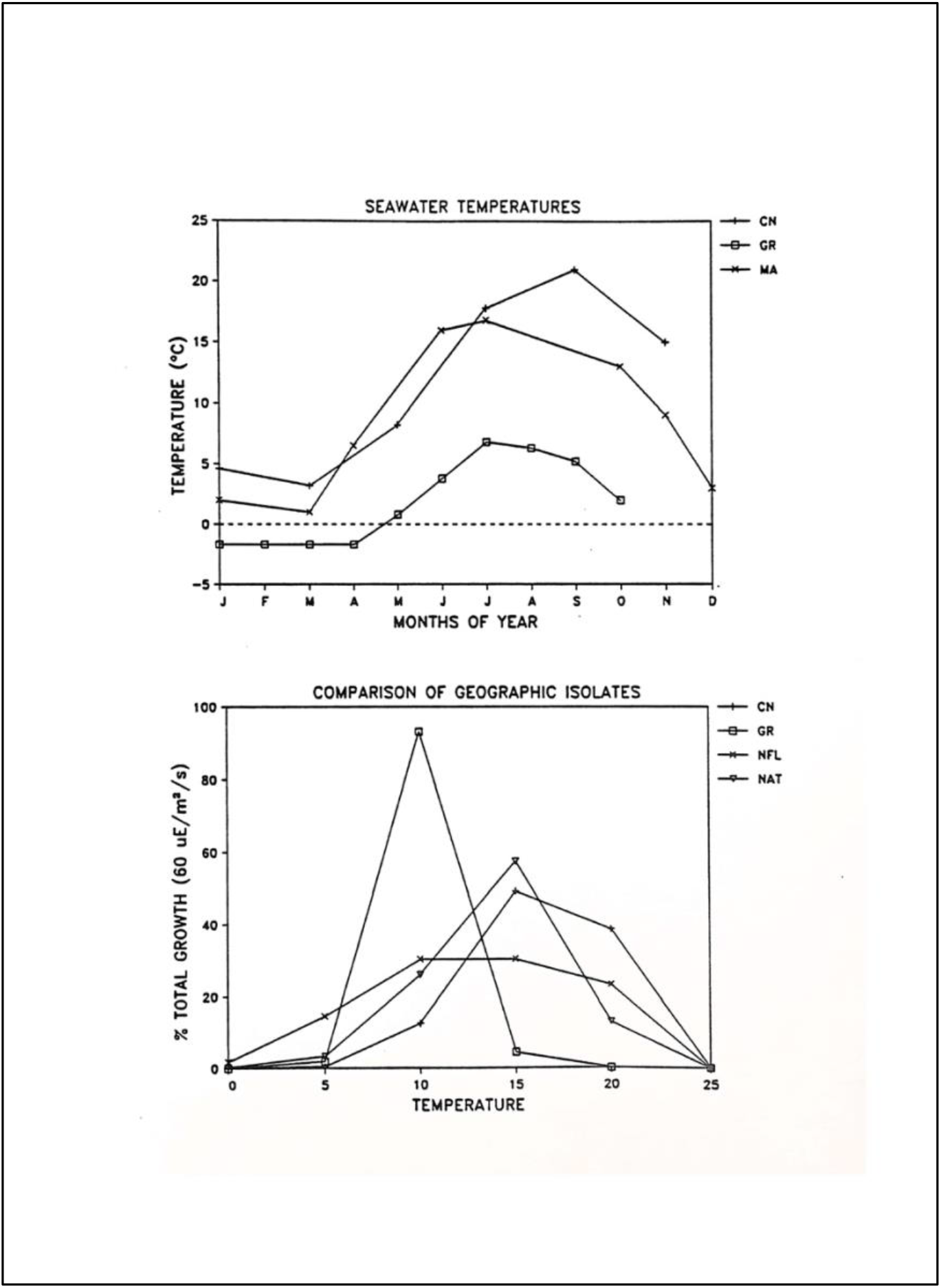
Monthly seawater temperatures and growth response to varying temperatures for geographic isolates of *Pilayella littoralis.* Seawater temperatures: Massachusetts data (MA) adapted from half-month average water temperatures taken at Northeastern University Marine Laboratory from January 1977 to June 1987 (pers. comm., C. Ellis). Connecticut data (CT) are adapted from Figure 4, page 10 of Northeast Utilities Environmental Lab (1987), representing averages from 1982 to 1986. Greenland data (GR) are for 1978 and from Table 2 of Anderson (1981). Growth response to varying temperatures; CN = Connecticut isolate. GR = Greenland isolate. NFL = average of two Nahant Bay, Massachusetts, free-living isolates. NAT = average of four Nahant Bay, Massachusetts, attached isolates.

**Figure 3.**
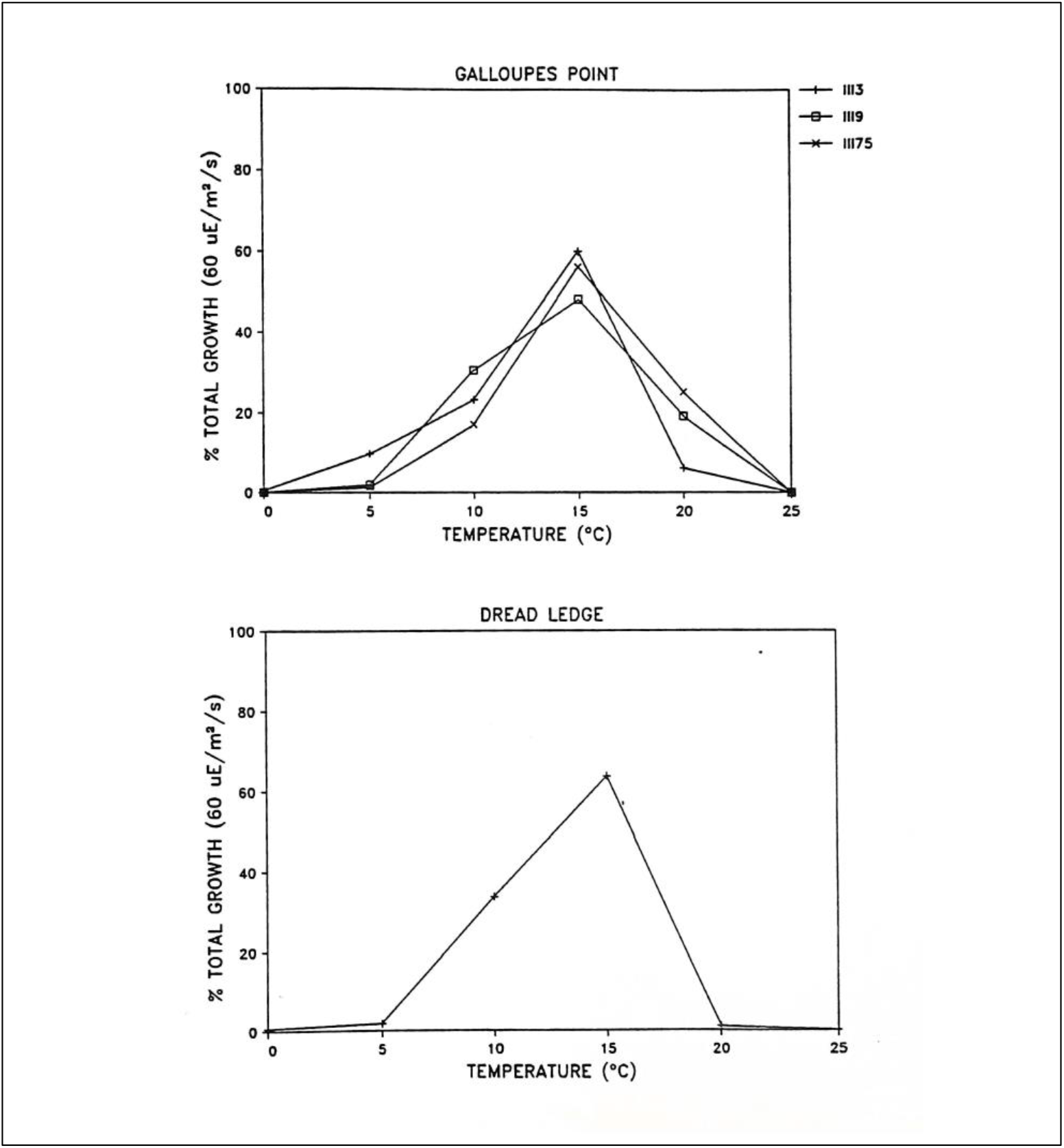
Growth responses of Nahant Bay attached *Pilayella littoralis* to varying temperatures. Epiphytic Galloupes Point algae include three 1984 summer isolates; III3 (run ten months after collection) and III9 (run 18 months after collection) under long days, and III75 (run 20 months after collection) under short days. Epilithic Dread Ledge algae is from a 1985 summer isolate; DL-98 (run after 3H months) under long days.

**Figure 4.**
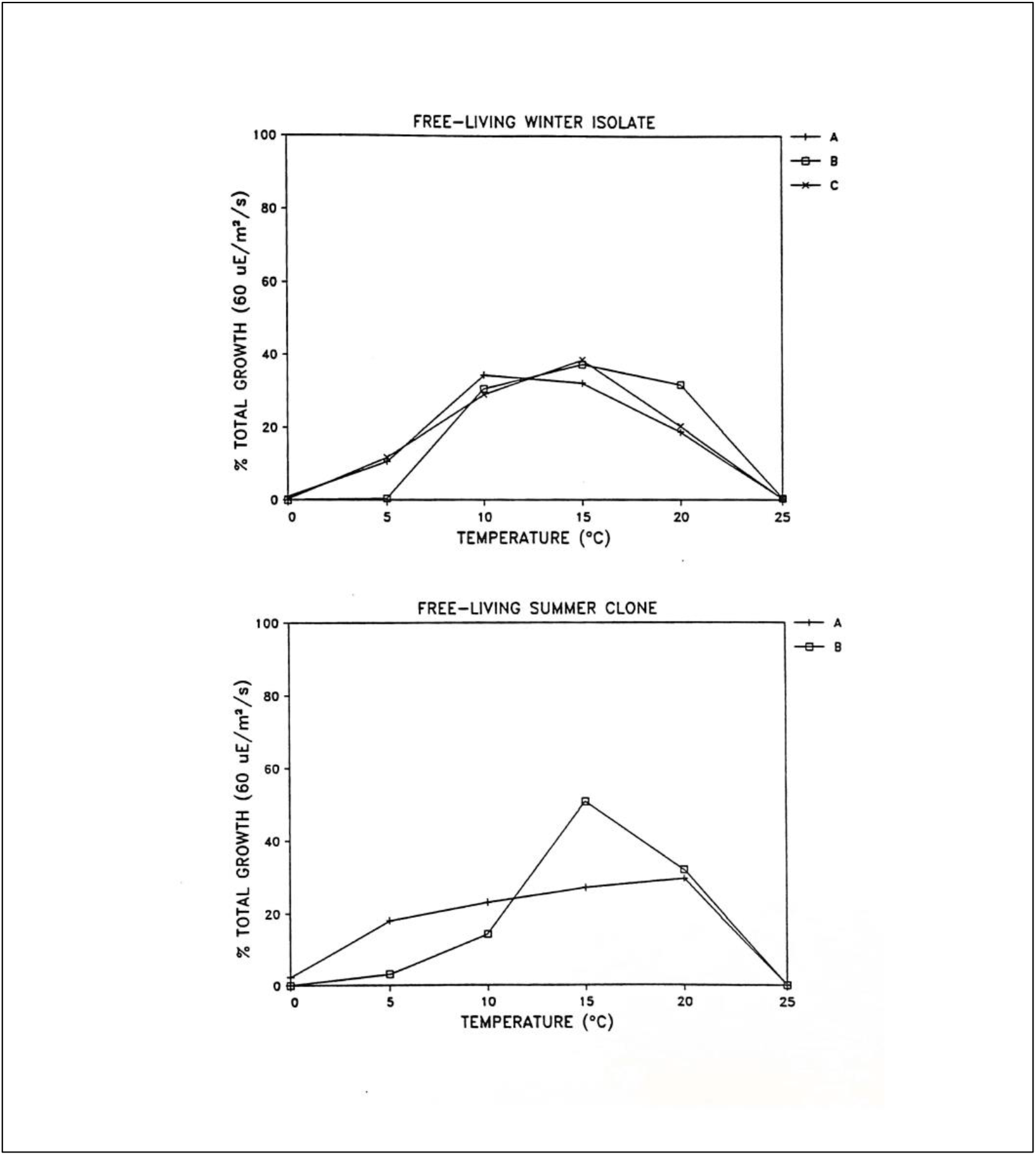
Growth responses of Nahant Bay free-living *Pilayella littoralis* to varying temperatures. The winter isolate 1143 (collected 2/12/84) was replicated three times; A was run 20 months after collection, B 24 months after collection, and C 34 months after collection. A and C were under long days, and B was under short days. The summer isolate III25 (collected 6/25/84) was replicated twice; A was run 12 months after collection, and B 41 months after collection, both under long days.

**Figure 5.**
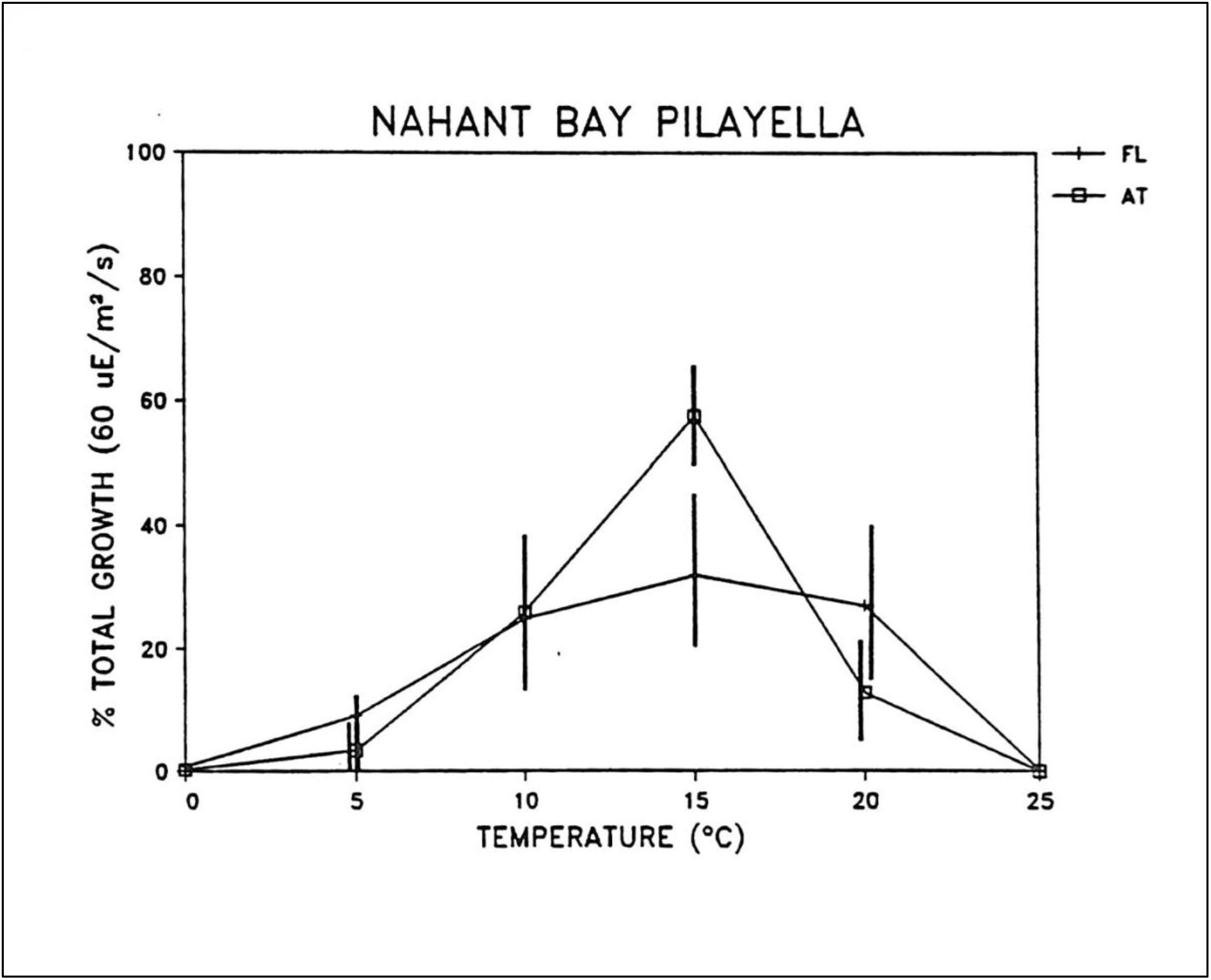
Average growth responses of Nahant Bay attached (N=4) and free-living *Pilayella littoralis* (N=2) to varying temperatures. Data for free-living algae (FL) include the average of three winter replicates and the average of two summer replicates. Data for the attached algae (AT) include three Galloupes Point isolates and one Dread Ledge isolate. Error bars are 95% confidence intervals.

ANOVA rejected equality of means within each temperature treatment for each algal form (significance level = 0.01). Scheffe’s Multiple Range Test demonstrated that growth at 15° C for attached algae was significantly greater than in other temperature treatments (significance level = 0.05). For free-living *P. littoralis,* a small sample size (n = 2) makes growth comparisons among treatments uncertain; even so, growth differences in 5°, 10°, 15°, and 20° C treatments were not statistically significant (Scheffe’s Multiple Range Test, significance level = 0.05). Figures 6 – 8 present photographic records of paired cross-gradient growth studies for Nahant Bay attached and free-living *P. littoralis*. Photographic negatives were accidentally destroyed in the first cross-gradient growth study, so these data are presented graphically in Figure 9.

**Figure 6.**
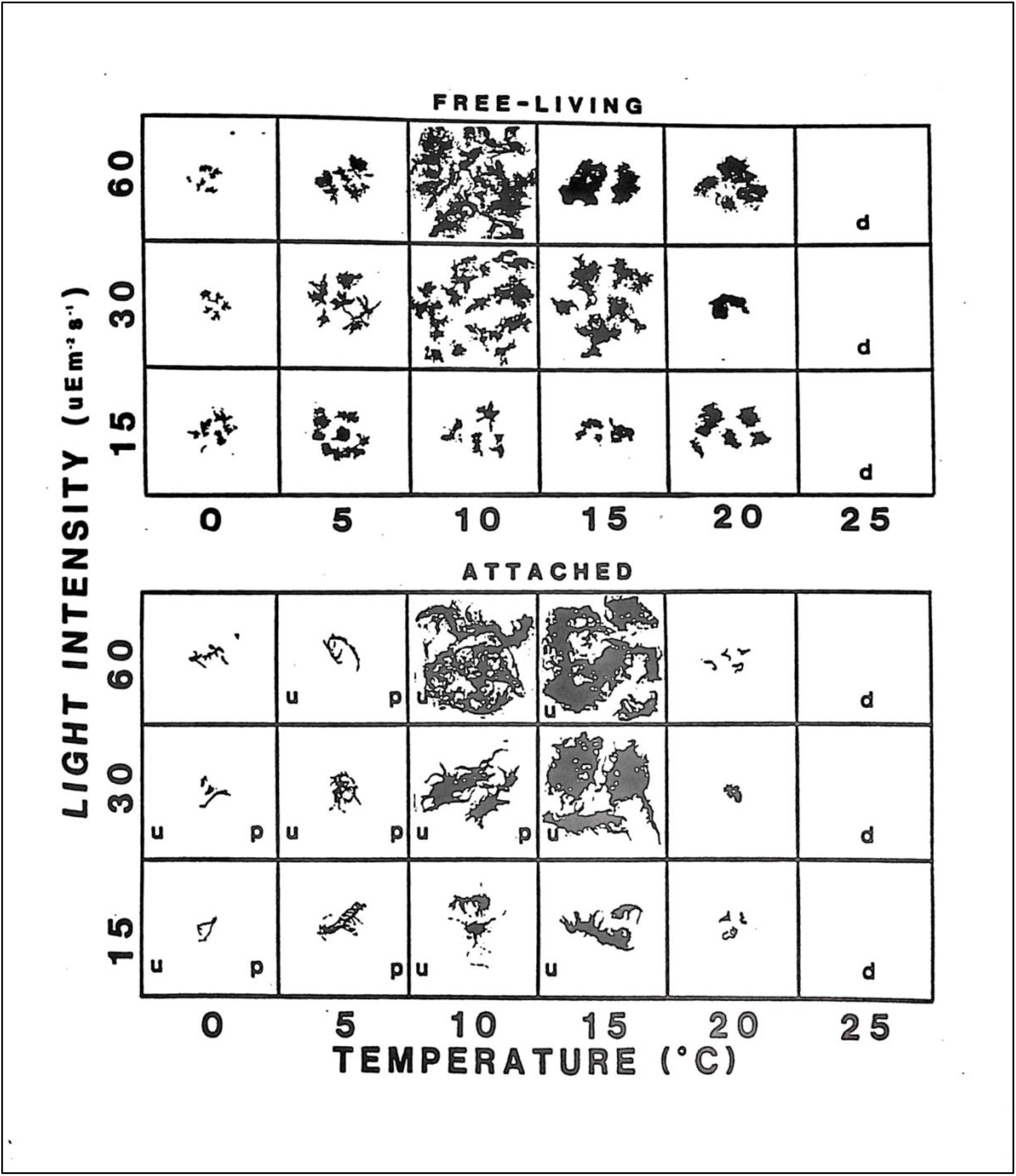
Growth responses of Nahant Bay, Massachusetts free-living (summer isolate) and attached (Dread Ledge isolate) *Pilayella littoralis* to varying light and temperature regimes. Duration of study 10/24/85 to 11/27/85 under long-day photoperiod (16:8, L:D). Legend: p = plurilocular reproductive organs, u = unilocular reproductive organs, s = survived duration of the experiment as tested by re-incubation at 12.5 °C, d = did not survive the experiment. Scale; sides of squares = 5 cm.

**Figure 7.**
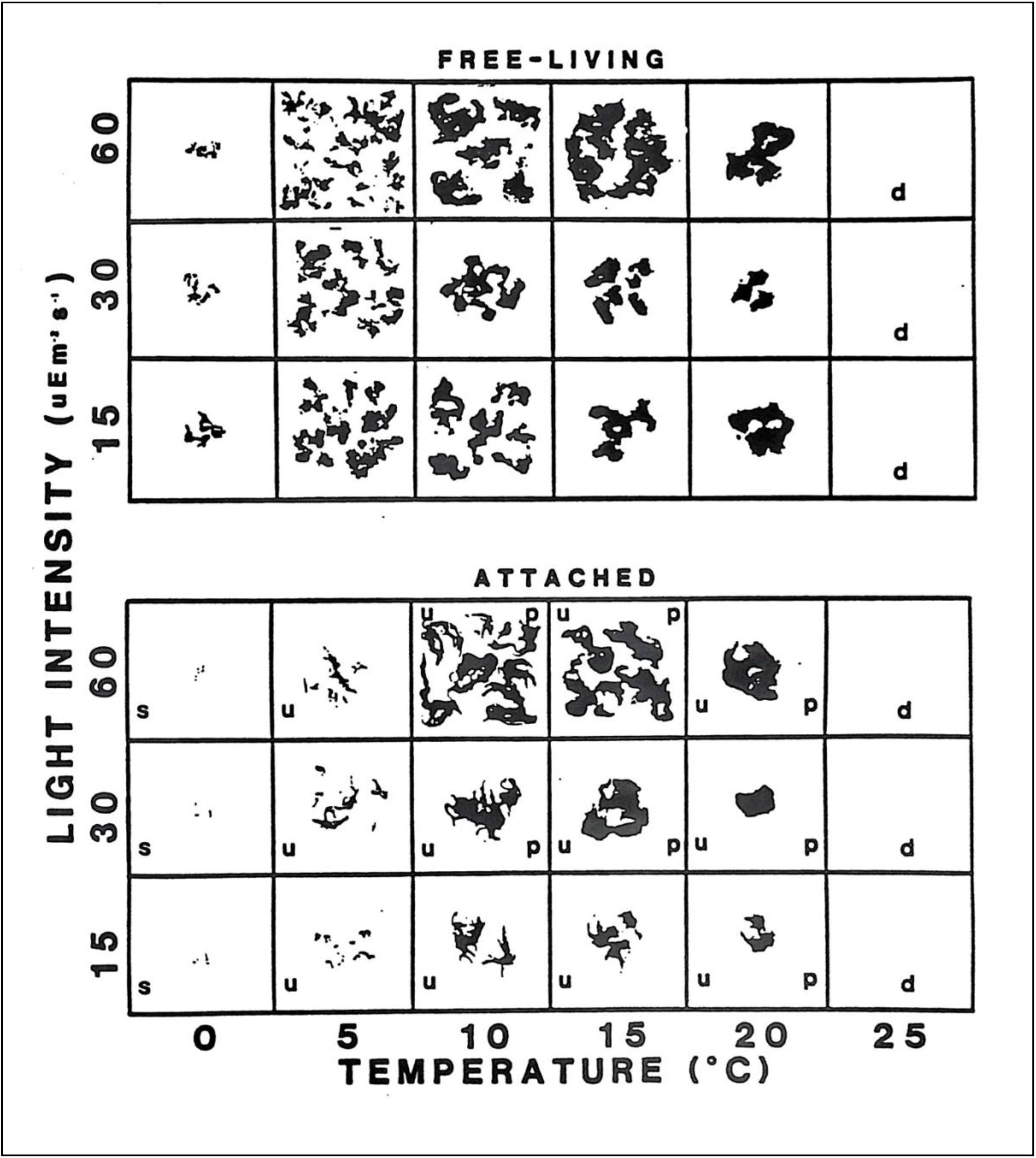
Growth responses of Nahant Bay, Massachusetts free-living (summer isolate) and attached (Galloupes Point, III9) *Pilayella littoralis* to varying light and temperature regimes, 12/26/85 to 1/29/86 under long-day photoperiod (16:8, L:D). Legend: p = plurilocular reproductive organs, u = unilocular reproductive organs, s = survived duration of the experiment as tested by re-incubation at 12.5 °C, d = did not survive the experiment. Scale; sides of squares = 5 cm.

**Figure 8.**
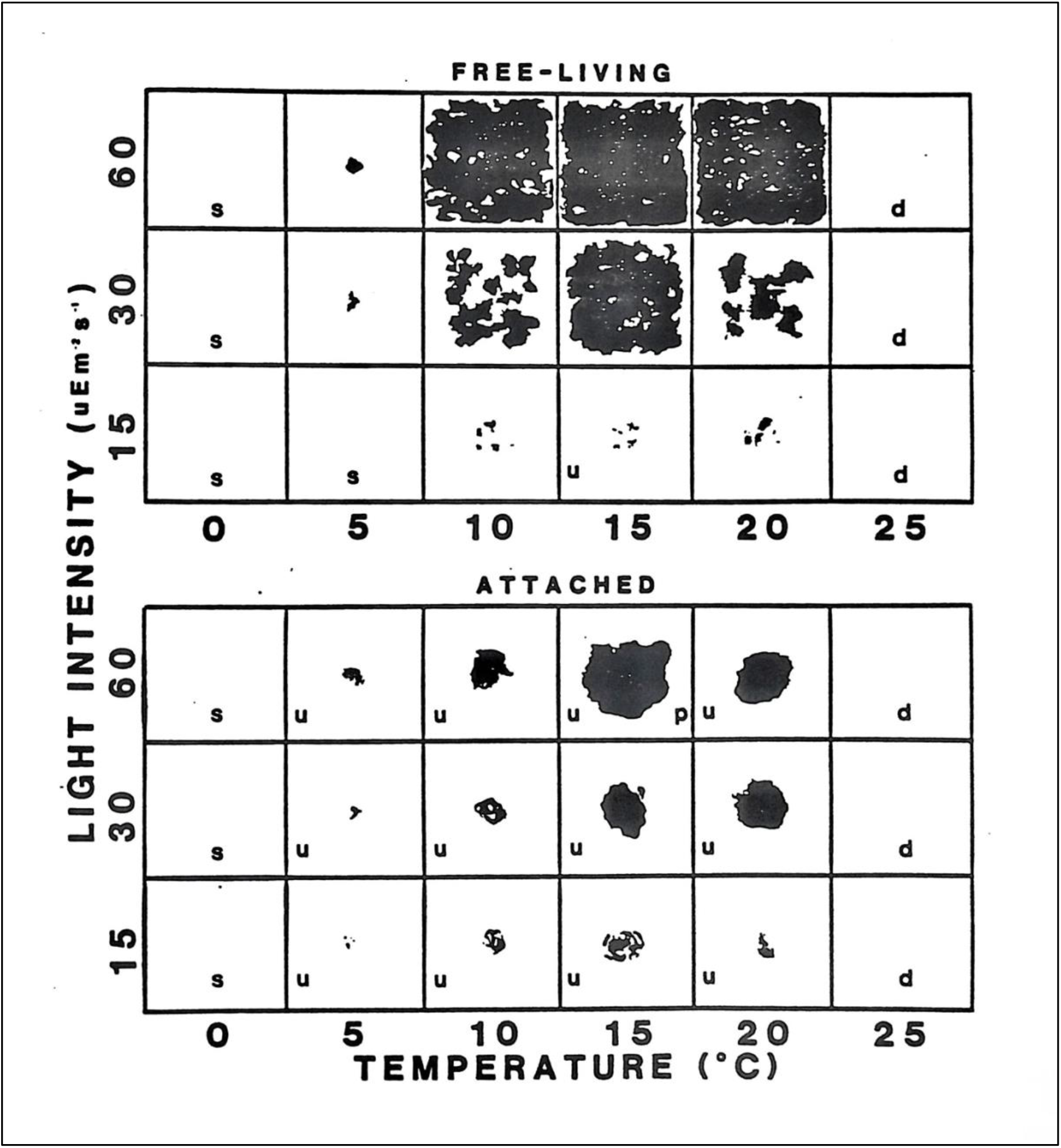
Growth responses of Nahant Bay, Massachusetts free-living (summer isolate) and attached (Galoupes Point, III75) *Pilayella littoralis* to varying light and temperature regimes, 2/11/86 to 3/27/86 under short-day photoperiod (8:16, L:D). Legend: p = plurilocular reproductive organs, u = unilocular reproductive organs, s = survived duration of the experiment as tested by reincubation at 12.5 °C, d = did not survive the experiment. Scale; sides of squares = 5 cm.

Growth responses were not the only difference between attached and free-living *Pilayella littoralis.* The general morphology of attached and free-living *P. littoralis* differed in all culture experiments. Attached plants exhibited cabled filaments, resulting in a clumped, twisted, or coarse appearance; pluriseriate filaments were uncommon. Free-living individuals infrequently produced cabled filaments and were extensively and loosely branched, ball-like in appearance, and often with pluriseriate axes. Clones of attached *P. littoralis* commonly produced unilocular reproductive structures and less frequently plurilocular reproductive organs. Free-living *P. littoralis,* except in one cross-gradient study, never produced reproductive structures of either type. Several unilocular reproductive organs were seen at 5 and 15° C, under short days, and at low light conditions. Otherwise, all free-living material in culture remained vegetative.

Growth responses of attached *Pilayella littoralis* collected from Disko Bay, west Greenland, and Connecticut, U.S.A, were distinctive from both Nahant Bay free-living and attached isolates. Maximum growth of the epilithic Greenland alga occurred at 10° C, with growth at higher and lower temperatures significantly less (Figure 10). Unilocular reproductive organs were produced under long-day conditions. Under short-day conditions the same clone produced both unilocular and plurilocular reproductive structures and exhibited a somewhat broader growth response curve than under long-day conditions (Figure 11). In contrast, the Connecticut clone of epiphytic *P. littoralis* grew well between 10 and 20° C but reproduced significantly only at 20° C (Figure 12). Plurilocular reproductive structures were produced under both long- and short-day conditions. Unilocular sporangia were only seen at 20° C under long days at high light conditions. The Connecticut isolate exhibited a shift in maximum growth response to 10° C under short-day compared to 20° C under long-day photoperiod (Figure 11).

**Figure 9.**
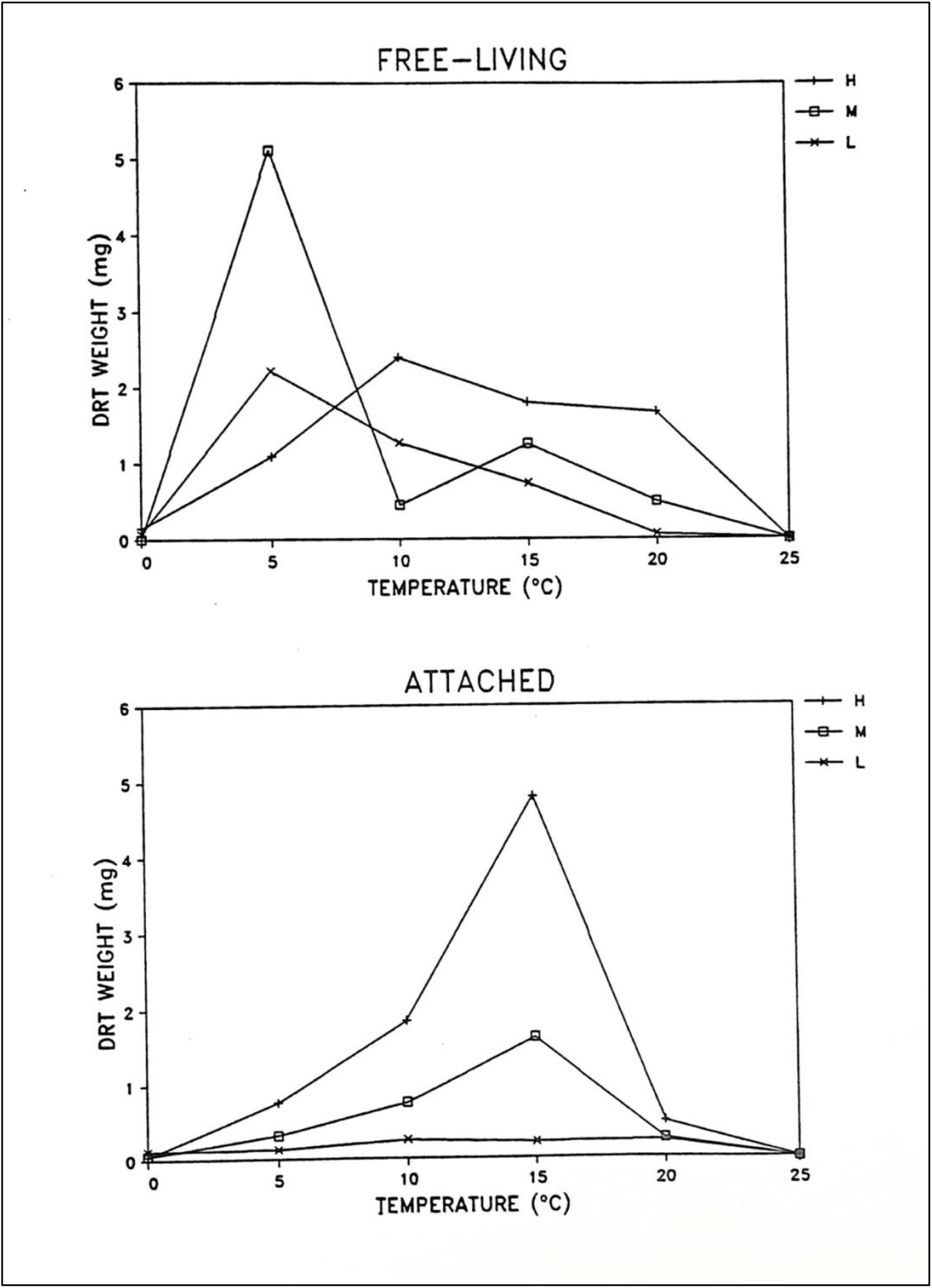
Growth responses of Nahant Bay, Massachusetts free-living (winter isolate) and attached (Galloupes Point, III3) *Pilayella littoralis* to varying light and temperature regimes, 5/3/85 to 6/21/85 under long-day photoperiod (16:8, L:D). Growth was measured as total biomass at the end of the experiment. H, M, and L are 60±5, 30±5, and 15±5 uE/m^2^/s respectively.

**Figure 10.**
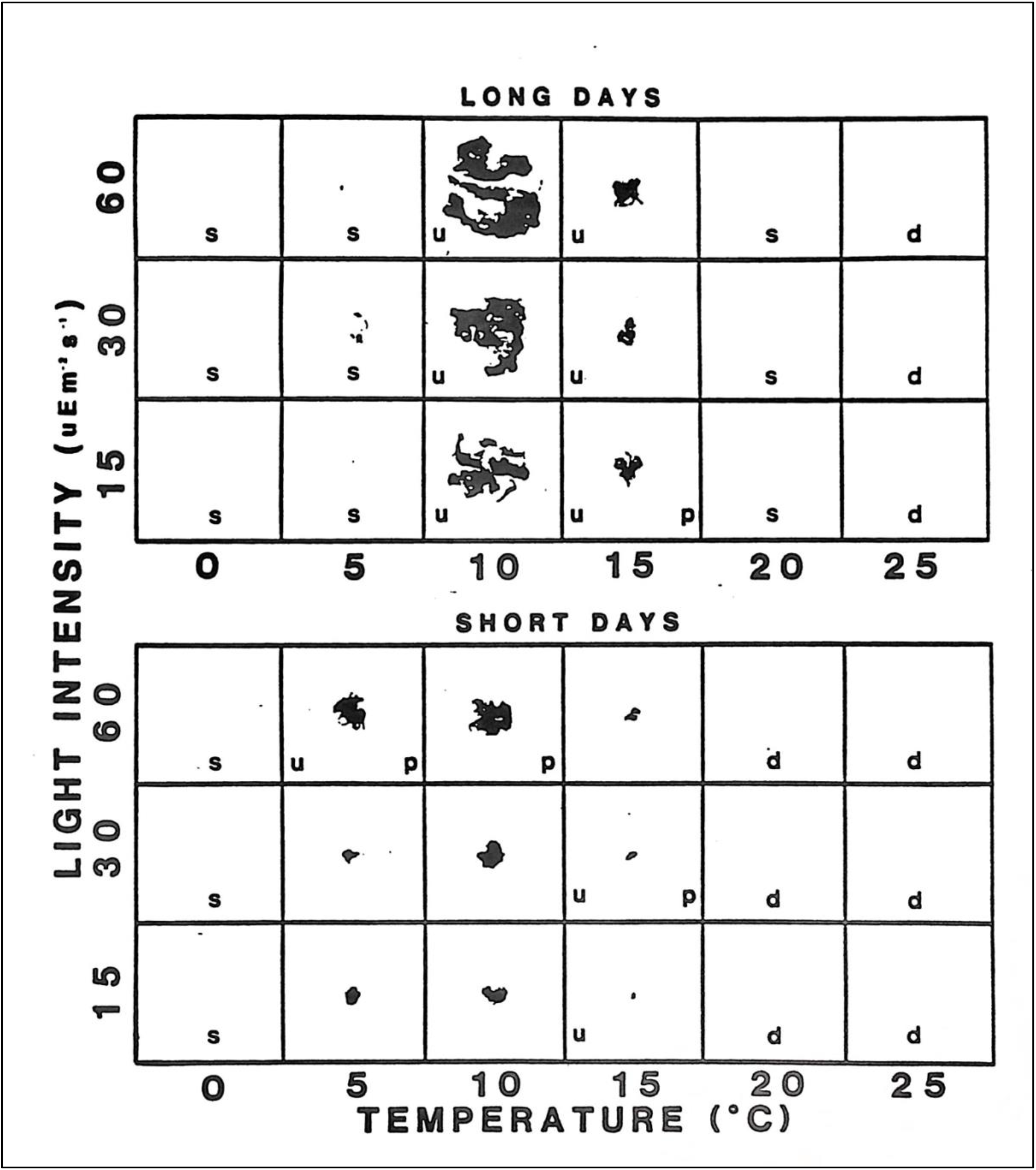
Growth responses of Disko Island, Greenland, attached *Pilayella littoralis* to varying light and temperature regimes, 5/11/87 to 6/15/87 under long-day photoperiod (16:8, L:D), and 10/2/87 to 10/29/87 under short-day photoperiod (8:16, L:D). Legend: p = plurilocular reproductive organs, u = unilocular reproductive organs, s = survived duration of the experiment as tested by re-incubation at 12.5 °C, d = did not survive the experiment. Scale; sides of squares = 5 cm.

**Figure 11.**
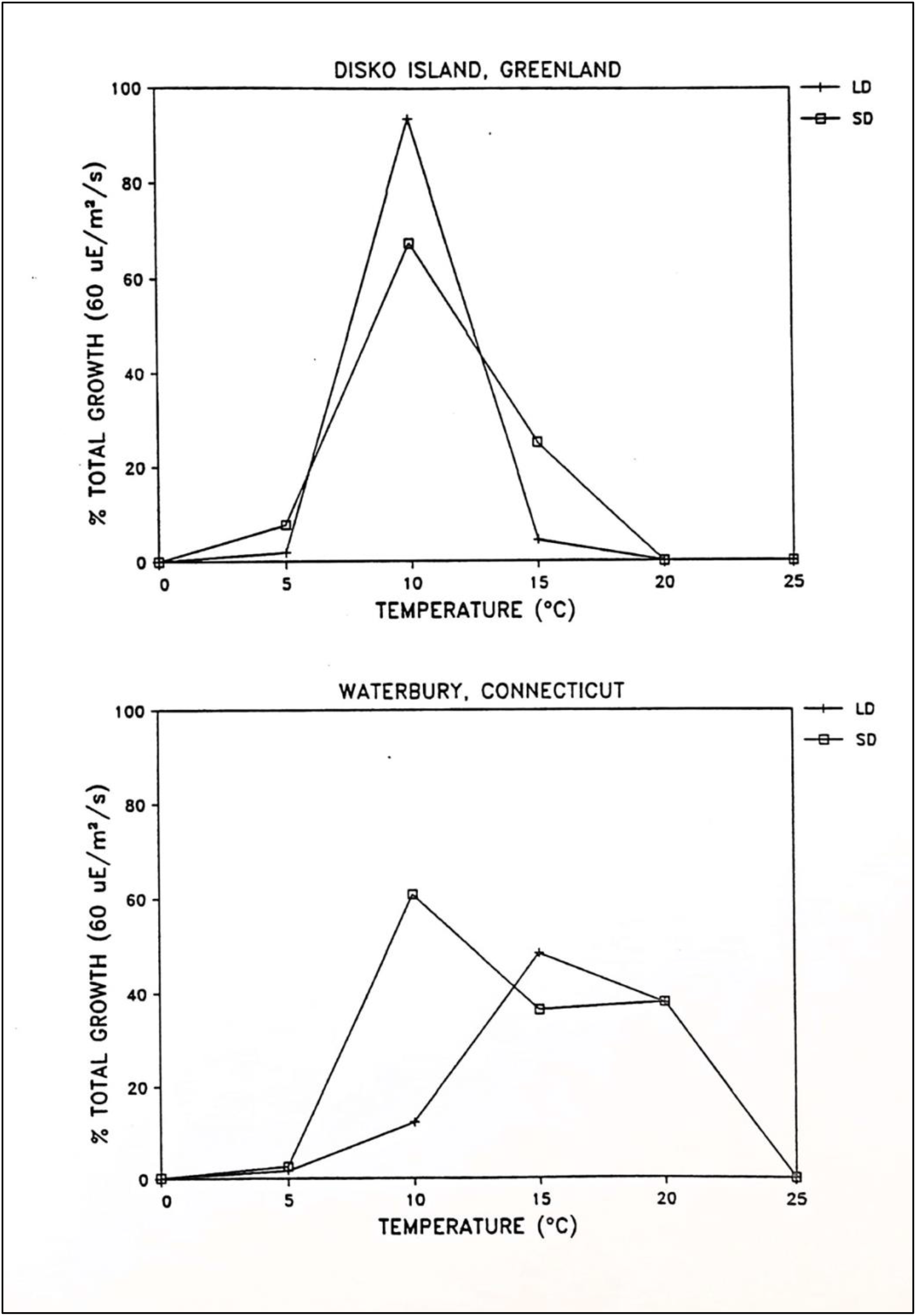
Growth responses of Disko Island, Greenland, and Connecticut attached *Pilayella littoralis* to varying light and temperature regimes. LD = long day photoperiod (16:8); SD = short day photoperiod (8:16); H, H, and L are 60±5, 30±5, and 15±5 uE/m^2^/s respectively.

**Figure 12.**
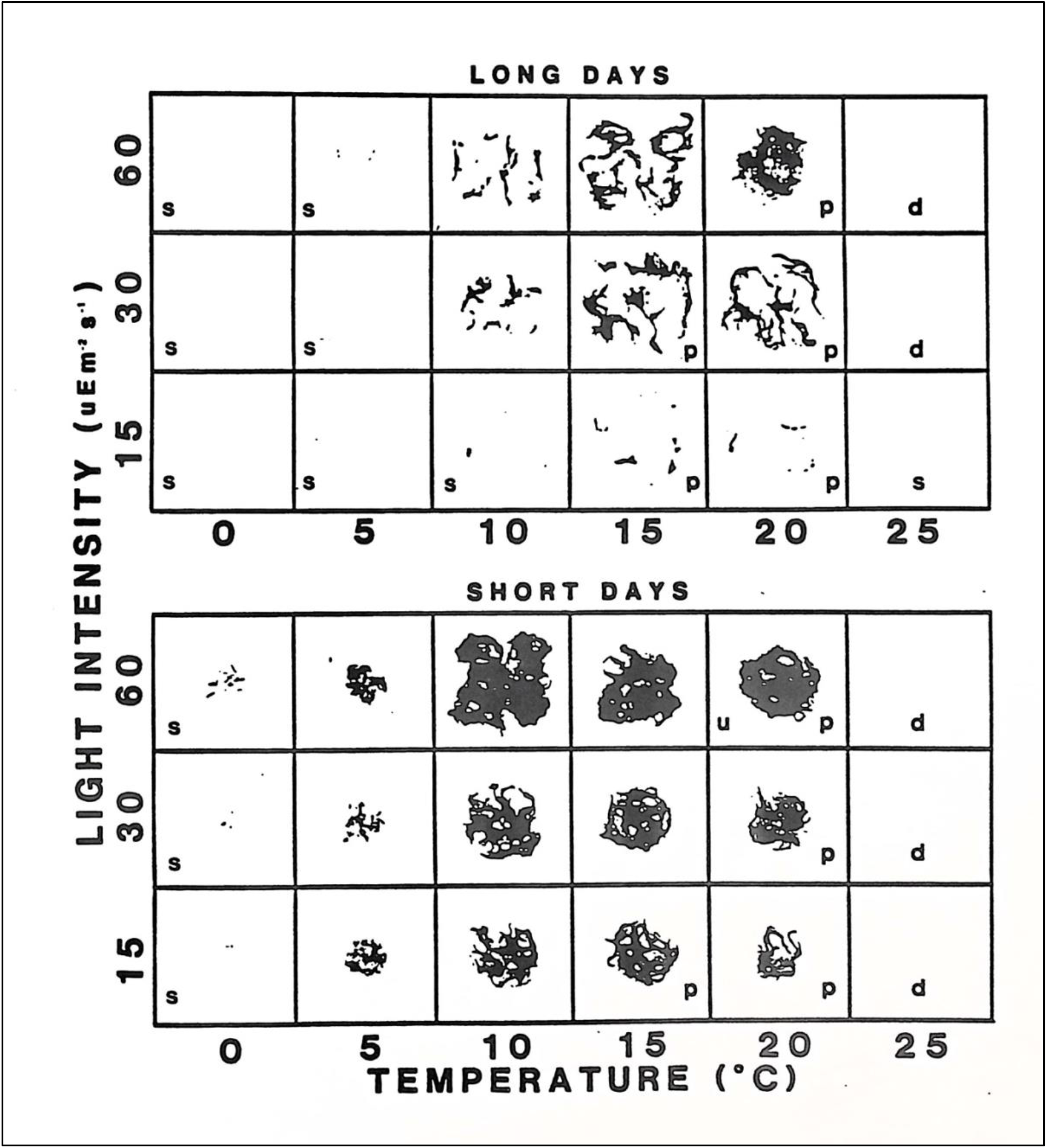
Growth responses of Connecticut attached *Pilayella littoralis* to varying light and temperature regimes. Duration of study 7/3/87 to 8/14/87 under long-day photoperiod (16:8), and 10/2/87 to 10/29/87 under short-day photoperiod (8:16). Legend: p = plurilocular reproductive organs, u = unilocular reproductive organs, s = survived duration of the experiment as tested by re-incubation at 12.5 °C, d = did not survive the experiment. Scale; sides of squares = 5 cm.

Under long days, growth for three isolates of epiphytic Nahant Bay attached *Pilayella littoralis* were similar (Figure 3). The Nahant Bay epilithic Dread Ledge isolate, however, had a relatively narrow growth response to temperature, growing significantly only between 10 - 15° C (Figure 3) compared with the epiphytic algae that grew well from 10 - 20° C, and moderately at 5° C.

Differences in growth response for isolates of Nahant Bay free-living *Pilayella littoralis* collected during winter and summer are not statistically significant (Figure 13, 95% confidence intervals for means at each temperature treatment overlap). Maximum growth in all treatments occurred under high irradiance, except in one instance. The greatest biomass developed at 5° C under medium irradiance for the summer free-living isolate after one year in culture (Figure 9). Since this experiment was run in duplicate with an attached isolate the difference in growth response between the two does not appear to be experimental error. Still, these results are anomalous as this isolate exhibited maximum growth at 15 - 20° C two years later under high irradiance (Figure 14). Also, considerably less biomass developed at 5° C than in the first replicate. Little difference in growth response was detected in three replicates of one free-living winter isolate over three years (Figure 4).

**Figure 13.**
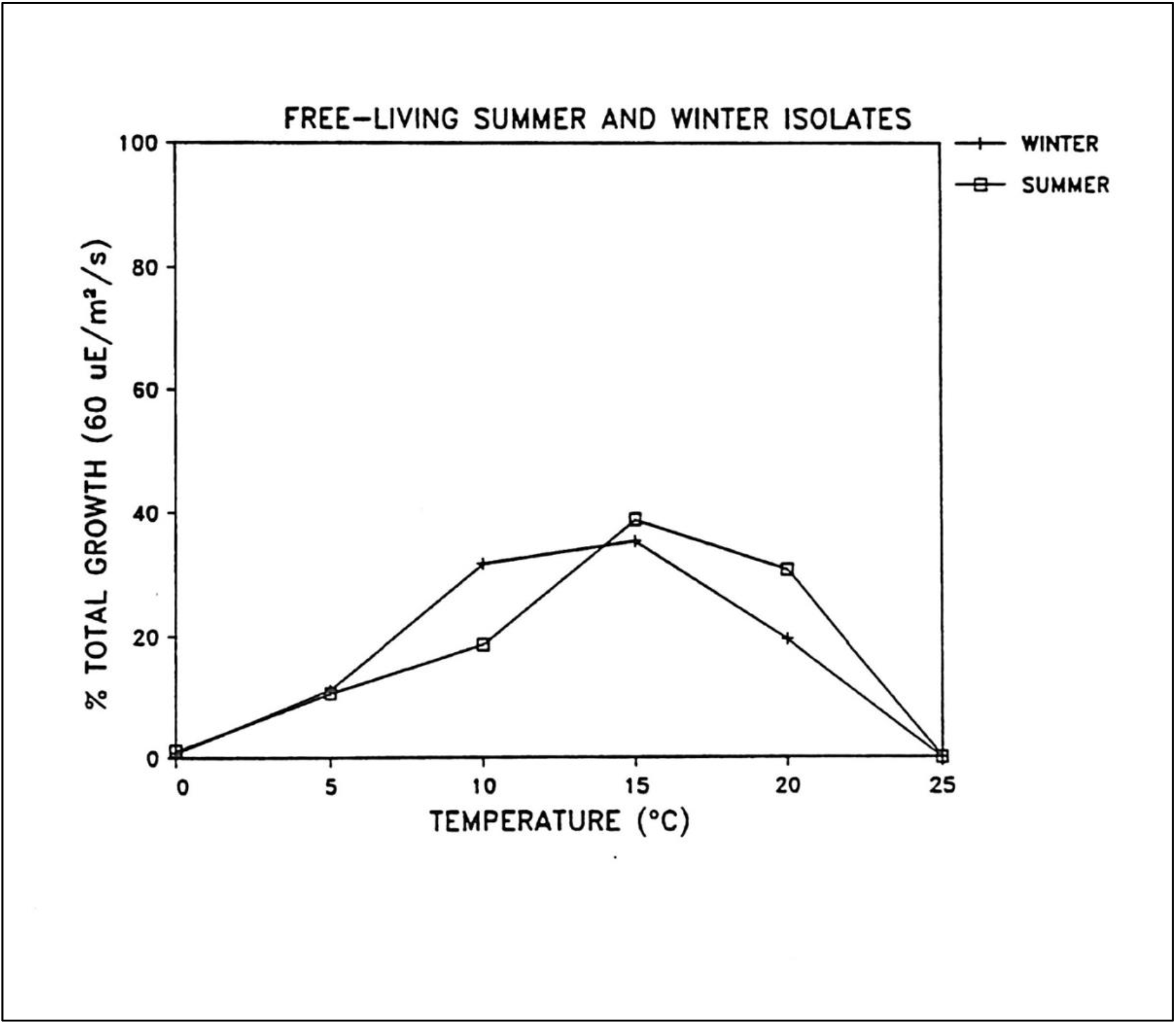
Average growth responses of Nahant Bay free-living *Pilayella littoralis* summer and winter isolates. The summer isolate III25 (collected 6/25/84) was replicated twice, and the winter isolate 1143 three times. Culture information as in Figure 4.

**Figure 14.**
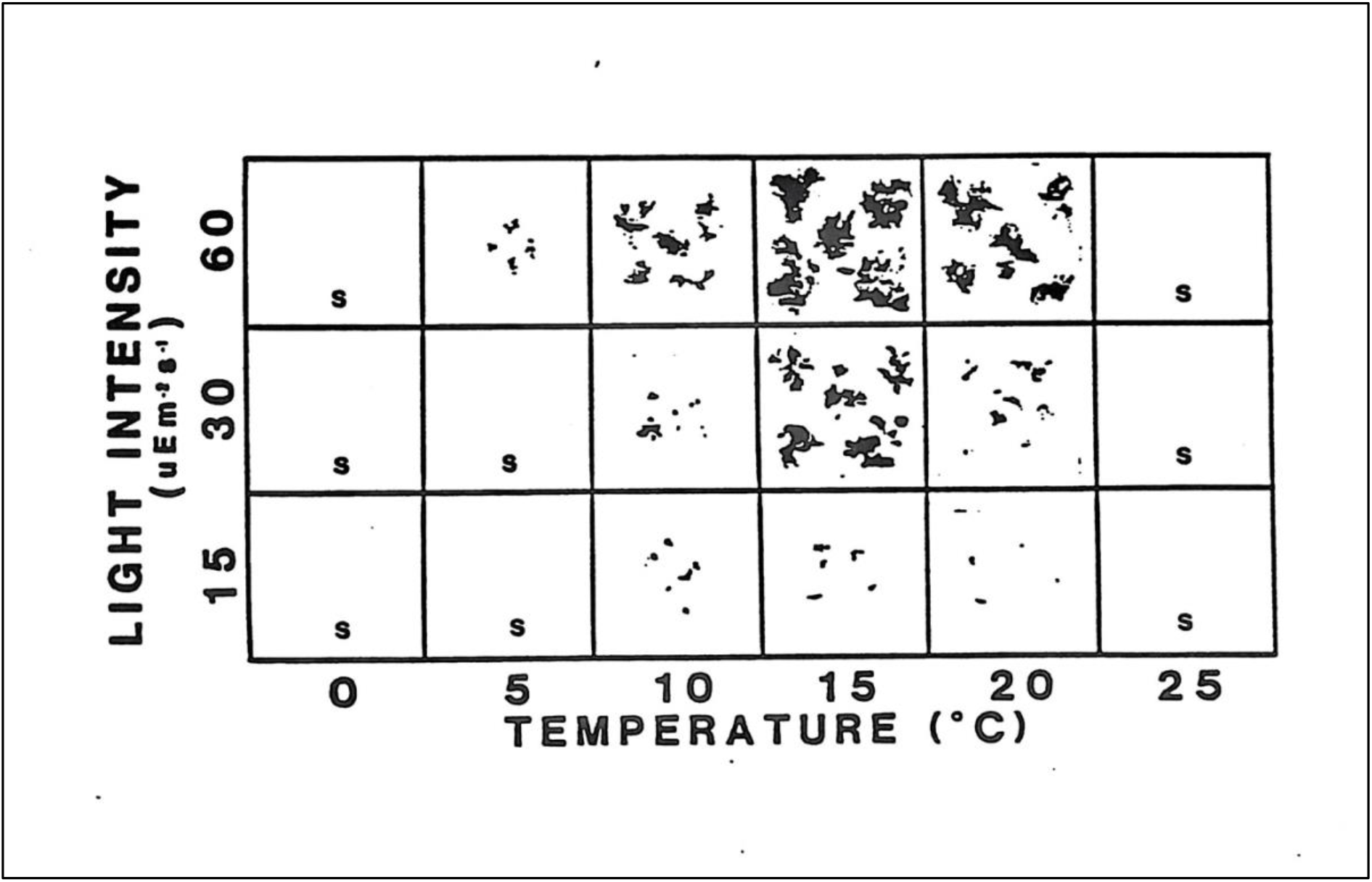
Growth responses of Nahant Bay, Massachusetts free-living *Pilayella littoralis* summer isolate (III25) to varying light and temperature regimes, 7/31/87 to 8/14/87. Compare results with the same isolate run 12 months after collection (Figure 4).

At high temperatures, near-lethal, both attached and free-living *Pilayella littoralis* exhibited atypical morphology. Attached plants from Nahant Bay were coarsely clumped and wiry in appearance at 20° C. Attached plants from Connecticut were also clumped at 20° C, but not as densely as those from Nahant. In the clumped morphology, axial regions were difficult to identify, and individual plants were impossible to discern. Growth of the Greenland isolate dropped off quickly at 15° C, and a clumped appearance was hardly apparent; filaments surviving 15° C were somewhat thickened, darkly pigmented, with short, spindly laterals. Coarse material was also seen in some free-living treatments, especially under long days, high irradiance and 20° C.

In cross-gradient experiments, the maximum survival of attached *Pilayella littoralis* collected from Dread Ledge, Nahant Bay, was near 20° C (algae were badly stressed at this temperature and did not survive 25° C, and between 20 and 25° C for attached algae collected from Galloupes Point (good growth at 20° C, but no growth at 25° C). The attached algae from Greenland survived one month at 20° C, but did not grow. Algae from Connecticut survived two weeks at 25° C but exhibited little growth and were stressed. Free-living *P. littoralis* survived all treatments at 20° C; only the summer clone survived two weeks at 25° C (Figure 14). Free-living *P. littoralis* that survived 25° C was severely stressed, and upon recovery exhibited a clumped cabled morphology similar to the attached form, except that reproductive organs were not produced. Apical cuttings from these plants subsequently produced a compact ball of unattached material, but no reproductive structures. These maximum survival temperatures are a rough approximation based on treatments at 20 and 25° C.

Growth rates (mg/day) of free-living and attached *Pilayella littoralis* from these cross-gradient studies are summarized in Tables 4-5. Specific growth rates (days to double biomass) are presented in Tables 6-7 and range from 2 to 17 days for free-living isolates and 4 to 17 days for attached isolates. Conclusions about interpopulation growth differences are complicated by the lack of replication and the utilization of growth media of two different nutrient concentrations (F/2 and F/4).

**Table 4.** Growth rate of free-living *Pilayella littoralis* in cross-gradient studies (mg/day). H, M, and L are 60 ±5, 30 ±5, and 15 ±5 uE/m^2^/s respectively; LD and SD are long-and short-day photoperiods (16:8, 8:16); s = no measurable growth, but survived the experiment based on re-incubation at 1.5 °C; d= dead, did not survive the experiment.

| Isolate ID | Temperature (°C)/Irradiance |  |  |  |  |  |  |
| --- | --- | --- | --- | --- | --- | --- | --- |
|  |  | 0 | 5 | 10 | 15 | 20 | 25 |
| I 143a<br>(LD) | H | .0017 | .0600 | .1486 | .1969 | .1037 | d |
|  | M | .0057 | .0371 | .0606 | .0614 | .0160 | d |
|  | L | .0020 | .0160 | .0369 | .0260 | .0489 | d |
| I 143b<br>(LD) | H | .0118 | .1226 | .4147 | .3456 | .2124 | d |
|  | M | .0050 | .1694 | .2921 | .3747 | .0709 | d |
|  | L | .0176 | .1300 | .1941 | .0888 |  | d |
| I 143c<br>(SD) | H | s | .0051 | .3487 | .4256 | .3615 | d |
|  | M | s | .0026 | .0667 | .2692 | .0667 | d |
|  | L | s | .0003 | .0013 | .0005 | .0051 | d |
| III 25a<br>(LD) | H | .0031 | .0222 | .0288 | .0367 | .0341 | d |
|  | M | .0002 | .1043 | .0092 | .0255 | .0100 | d |
|  | L | .0016 | .0453 | .0259 | .0149 | .0112 | d |
| III 25b<br>(LD) | H | .0007 | .0107 | .0480 | .1720 | .1080 | s |
|  | M | .0007 | .0033 | .0193 | .1073 | .0433 | s |
|  | L | .0007 | .0047 | .0100 | .0127 | .0087 | s |

**Table 5.** Growth rate of attached *Pilayella littoralis* in cross-gradient studies (mg/day). H, M, and L are 60 ±5, 30 ±5, and 15 ±5 uE/m^2^/s respectively; LD and SD are long- and short-day photoperiods (16:8, 8:16); s = no measurable growth, but survived the experiment based on re-incubation at 1.5 °C; d= dead, did not survive the experiment.

| Isolate | Temperature (°C)/Irradiance |  |  |  |  |  |
| --- | --- | --- | --- | --- | --- | --- |
|  | 0 | 5 | 10 | 15 | 20 | 25 |
| M'stone, CN<br>(LD) | H s | .0087 | .0280 | .1113 | .0873 | d |
|  | M s | .0040 | .0180 | .0713 | .0527 | d |
|  | L s | s | .0187 | .0187 | .0113 | s |
| M'stone, CN<br>(SD) | H .0007 | .0122 | .2759 | .1648 | .1719 | d |
|  | M .0011 | .0044 | .0837 | .0907 | .0559 | d |
|  | L s | .0059 | .0537 | .0589 | .0222 | d |
| Greenland<br>(LD) | H s | .0087 | .4373 | .0207 | s | d |
|  | M s | .0087 | .2600 | .0033 | s | d |
|  | L s | .0013 | .1293 | .0113 | s | d |
| Greenland<br>(SD) | H s | .0053 | .0967 | .0360 | s | d |
|  | M s | .0060 | .0313 | .0087 | s | d |
|  | L s | .0047 | .0127 | .0067 | s | d |
| III 9<br>(LD) | H .0012 | .0126 | .2097 | .3306 | .1318 | d |
|  | M .0006 | .0147 | .0823 | .1285 | .0412 | d |
|  | L .0041 | .0088 | .0262 | .0385 | .0162 | d |
| III 3<br>(LD) | H .0010 | .0155 | .0376 | .0978 | .0098 | d |
|  | M .0012 | .0065 | .0153 | .0329 | .0051 | d |
|  | L .0024 | .0027 | .0051 | .0043 | .0047 | d |
| III 75<br>(SD) | H s | .0068 | .0864 | .2841 | .1273 | d |
|  | M s | .0045 | .0102 | .0364 | .0909 | d |
|  | L s | s | .0007 | .0042 | .0030 | d |
| DL 98<br>(LD) | H .0044 | .0088 | .3206 | .6265 | .0074 | d |
|  | M .0059 | .0115 | .2238 | .4121 | .0088 | d |
|  | L .0026 | .0121 | .0638 | .1138 | .0026 | d |

**Table 6.** Specific growth rate of free-living *Pilayella littoralis* in cross-gradient growth studies. Data represented as days to double biomass. H, M, and L are 60 ±5, and 15 ±5 uE/m^2^/s respectively; LD and SD are long- and short-days (16:8, 8:16); F/2 and F/ were growth media; s = no measurable growth, but survived the experiment based on re-incubation at 12.5 °C; d = dead, did not survive the experiment.

| Isolate | Temperature (°C)/Irradiance |  |  |  |  |  |
| --- | --- | --- | --- | --- | --- | --- |
|  | 0 | 5 | 10 | 15 | 20 | 25 |
| I 143a | H 12.5 | 5.2 | 4.6 | 4.4 | 4.8 | d |
| (LD) | M 8.1 | 5.6 | 5.2 | 5.2 | 6.5 | d |
| (F/2) | L 10.8 | 6.5 | 5.6 | 6.0 | 5.3 | d |
| I 143b | H 6.9 | 4.7 | 4.0 | 4.1 | 4.4 | d |
| (LD) | M 8.4 | 4.5 | 4.2 | 4.1 | 5.1 | d |
| (F/2) | L 6.4 | 4.7 | 4.4 | 4.9 | 4.3 | d |
| I 143c | H s | 9.5 | 4.8 | 4.7 | 4.7 | d |
| (SD) | M s | 11.4 | 5.9 | 4.9 | 5.9 | d |
| (F/4) | L s | s | 14.3 | 21.3 | 9.5 | d |
| III 25a | H s | 3.8 | 2.7 | 2.2 | 2.4 | d |
| (LD) | M s | 5.3 | 3.3 | 2.4 | 2.8 | d |
| (F/2) | L s | 4.8 | 3.8 | 3.6 | 4.0 | d |
| III 25b | H 12.8 | 8.5 | 8.1 | 7.9 | 7.8 | s |
| (LD) | M s | 7.1 | 10.0 | 8.3 | 9.8 | s |
| (F/4) | L 15.2 | 7.6 | 8.31 | 9.1 | 16.6 | s |

**Table 7.** Specific growth rate of attached *Pilayella littoralis* in cross-gradient growth studies. Data represented as days to double biomass. H, M, and L are 60 ±5, and 15 ±5 uE/m^2^/s respectively; LD and SD are long- and short-days (16:8, 8:16); F/2 and F/4 were growth media; s = no measurable growth, but survived the experiment based on re-incubation at 12.5 °C; d = dead, did not survive the experiment.

| Isolate | Temperature (°C)/Irradiance |  |  |  |  |  |
| --- | --- | --- | --- | --- | --- | --- |
|  | 0 | 5 | 10 | 15 | 20 | 25 |
| M' stone,<br>(LD)<br>(F/4) | H s<br>M s<br>L s | 12.5<br>5.0<br>s | 3.0<br>3.3<br>3.3 | 2.4<br>2.6<br>3.3 | 2.5<br>2.7<br>3.7 | d<br>d<br>s |
| M' stone,<br>(SD)<br>(F/4) | H 14.3<br>M 11.8<br>L s | 5.7<br>7.3<br>6.8 | 3.4<br>4.1<br>4.4 | 3.7<br>4.0<br>4.3 | 3.7<br>4.3<br>5.1 | d<br>d<br>d |
| Greenland<br>(LD)<br>(F/4) | H s<br>M s<br>L s | 9.3<br>9.3<br>18.5 | 4.5<br>4.9<br>5.4 | 7.5<br>12.5<br>8.7 | s<br>s<br>s | d<br>d<br>d |
| Greenland<br>(SD)<br>(F/4) | H s<br>M s<br>L s | 8.2<br>7.9<br>8.5 | 4.4<br>5.3<br>6.5 | 5.2<br>7.1<br>7.7 | s<br>s<br>s | d<br>d<br>d |
| III 9<br>(LD)<br>(F/2) | H 13.5<br>M 18.5<br>L 9.1 | 7.0<br>6.8<br>7.6 | 4.5<br>5.1<br>6.1 | 4.3<br>4.8<br>5.7 | 4.8<br>5.7<br>6.7 | d<br>d<br>d |
| III 3<br>(LD)<br>(F/2) | H 17.8<br>M 16.6<br>L 13.5 | 9.1<br>10.8<br>13.7 | 7.8<br>9.1<br>11.4 | 6.8<br>7.9<br>11.8 | 9.8<br>11.4<br>11.5 | d<br>d<br>d |
| III 75<br>(SD)<br>(F/4) | H s<br>M s<br>L s | 8.7<br>9.5<br>s | 5.6<br>8.0<br>17.2 | 4.8<br>6.4<br>9.6 | 5.3<br>5.5<br>10.6 | d<br>d<br>d |
| DL 98<br>(LD)<br>(F/2) | H 8.2<br>M 8.1<br>L 10.0 | 7.4<br>7.0<br>6.9 | 4.1<br>4.3<br>5.2 | 3.8<br>4.0<br>4.7 | 7.7<br>7.4<br>10.0 | d<br>d<br>d |

### Effects of environment on morphology

The form and characteristics of free-living and attached *Pilayella littoralis* could not be transformed interchangeably *in vitro* (Figure 15). Cultures of attached plants bubbled continuously for eight months produced unilocular and plurilocular reproductive organs throughout the study, and never developed the nonpolar ball morphology of the free-living form. When apical cuttings approximately 1 mm long of attached unialgal material (epiphytic isolate from Galloupes Point, Massachusetts) were used as inocula, germlings subsequently developed on the bottom and sides of culture vessels in all treatments. Many of these attached germlings eventually detached and tumbled freely in culture. However, the morphology of these unattached individuals never resembled the morphology of free-living *P. littoralis.* Subcultures started using a variety of attached forms all grew and reproduced. However, none developed a form that resembled a typical “ball” of free-living *P. littoralis*.

**Figure 15.**
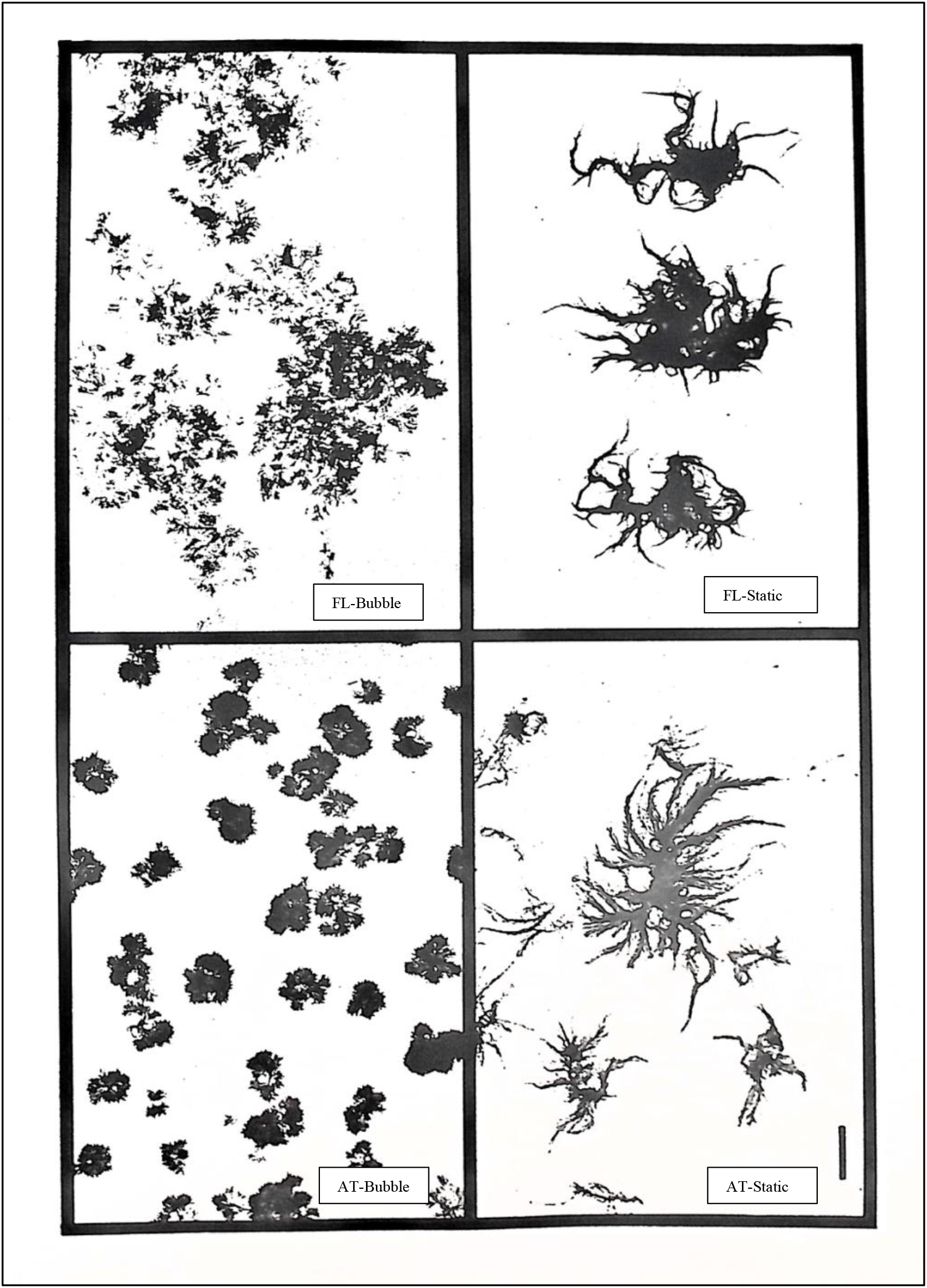
Growth responses of Nahant Bay free-living and attached *Pilayella littoralis* in bubbled and static cultures. Top; free-living isolate from bubble culture (left FL-Bubble) and static culture (right FL-Static). Bottom; attached isolate from bubbled (left AT-Bubble) and static culture (right AT-Static). Scale bar = 1 cm.

For example, motile cells from unilocular reproductive organs produced germlings attached to glass that subsequently became detached and grew free-living. These unattached individuals had a dense central region from which filaments originated. The growth of filaments was polar with respect to the dense central region, and the cabling of axes was common. The dense central region appeared to develop from the attachment pad of germlings and consisted initially of interwoven prostrate filaments and rhizoids. This region eventually became compact and was a nucleus for the developing individual. These compact ball-like structures also formed in cultures with inocula that included thickened axial regions and clumps that developed due to precocious germination within catenate unilocular sporangia.

Free-living *Pilayella littoralis* could not be induced to develop a form resembling the attached morphology. Bubbled cultures produced typical balls and static cultures produced large, delicate, loosely branched individuals without polarity (Figure 15). Further, free-living *P. littoralis* cultured in Pyrex screw-top 20 ml tubes for over four years, under static conditions with minimal maintenance, sometimes re-attached to the glass by rhizoids, and a slightly cabled less delicate form developed. However, these cultures generally do not resemble the attached form and revert to the typical ball form when cultured in larger vessels. For example, of 33 free-living isolates grown in 20 ml tubes for four years, two produced plurilocular reproductive organs, and three produced unilocular reproductive organs exclusively. In contrast, 22 attached isolates had some sort of reproductive organ; 19 had plurilocs, 3 had unilocs, and one had both plurilocs and unilocs.

Attached *Pilayella littoralis* collected from Nahant Bay, and grown in screw-top tubes as above, produced vegetative fragments that often grew unattached to glass. Motile reproductive cells produced by attached algae growing in culture tubes usually germinated after attaching to glass and infrequently at the interface between culture media and air.

### Electrophoresis

Of 39 enzymes assayed, six exhibited activity that could be resolved for analysis. The enzymes IDH, ALD, SKDH, SOD, and PRO (a general protein stain) were all monomorphic, with attached and free-living *Pilayella littoralis* sharing the same alleles (Table 8). The enzymes PGI and PGM exhibited polymorphisms in both forms of the species. However, banding patterns for PGM were not reproducible. As a result, no genetic interpretation could be made based on the PGM results. For PGI, two alleles were detected among attached and free-living *P. littoralis* with no individuals heterozygous (Figure 16 - 17). Few individuals were polymorphic. One of the 38 free-living isolates could be distinguished based on its PGI genotype (Figure 17); two of the 30 attached isolates shared this same allele with the free-living alga.

**Figure 16.**
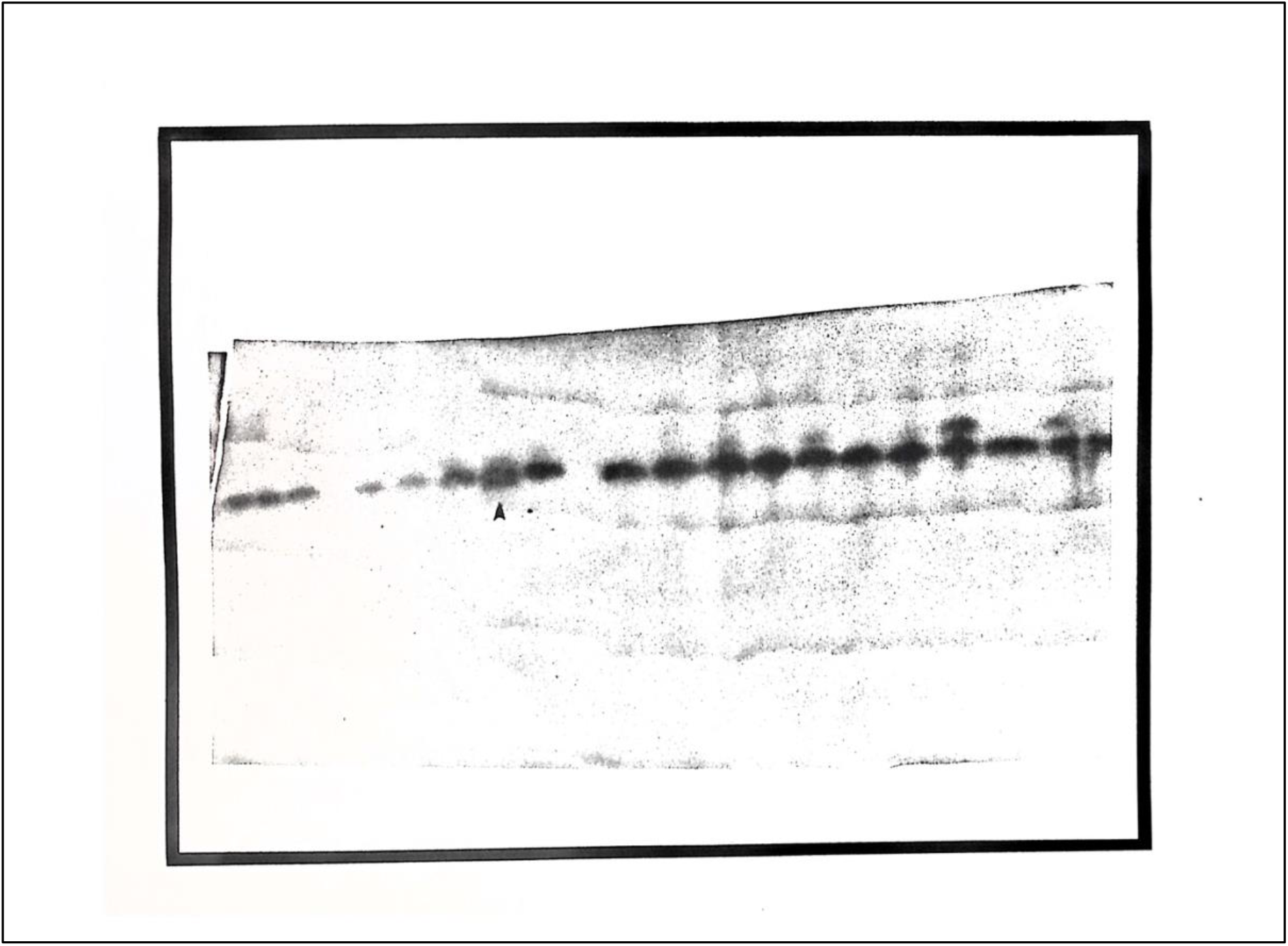
Isozyme variation among attached and free-living populations of Nahant Bay *Pilayella littoralis:* PGI enzyme assay. Bands 1-5; attached Dread Ledge isolates. Bands 6-10; attached Galloupes Point isolates. Bands 11-25; free-living winter isolates (four bands not visible at end of gel). Note the double bands in an attached Galloupes Point isolate (indicated by arrow). Double bands in other isolates are artifacts.

**Figure 17.**
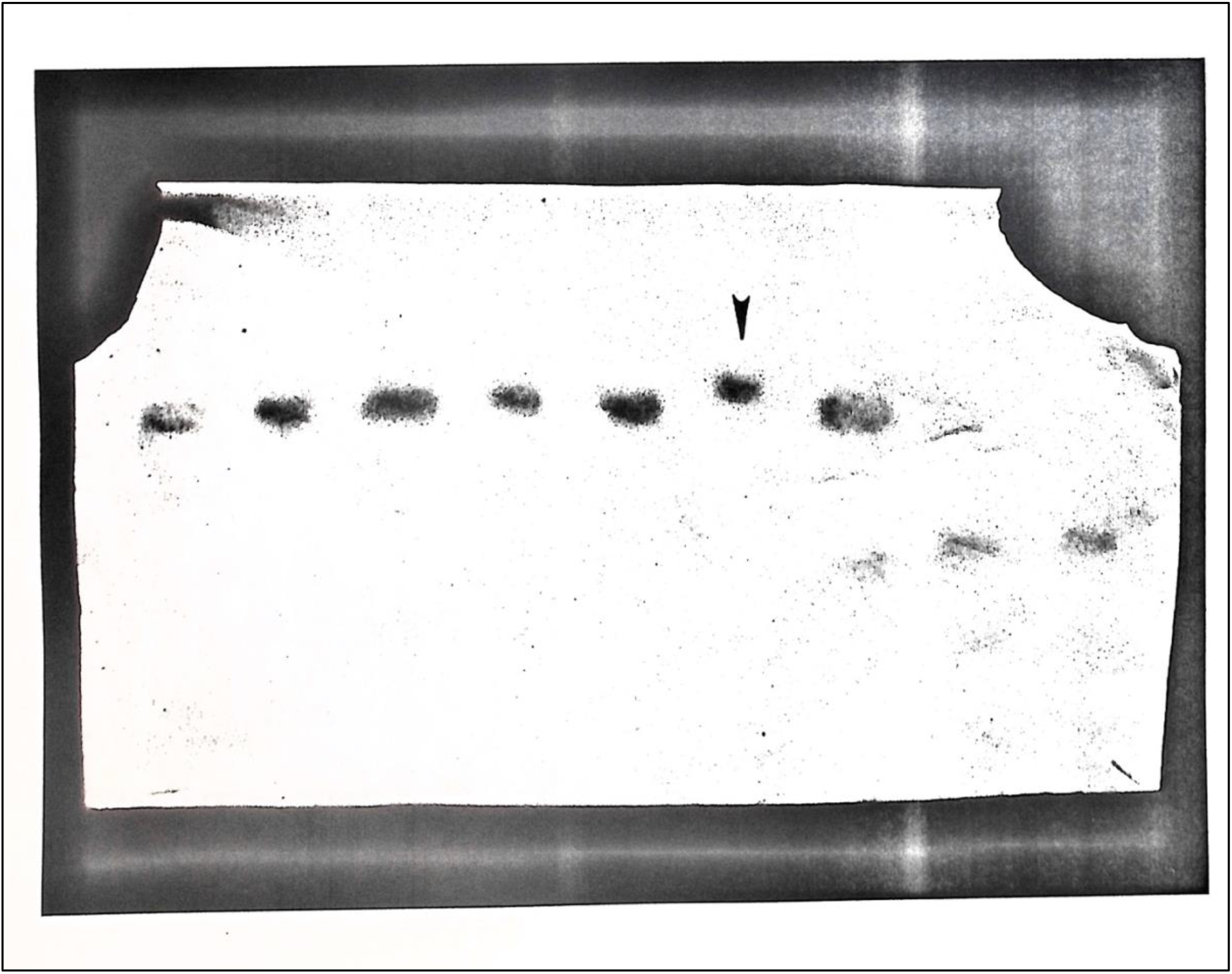
Isozyme variation among Nahant Bay free-living *Pilayella littoralis:* PGI enzyme assay. Arrow indicates an alternate allele in a summer isolate.

**Table 8.** Enzymes used in starch gel electrophoresis of *Pilayella littoralis*. Staining follows the recipes of Brewer (1970), Shaw and Prasad (1970), Harris and Hopkinson (1976), and Soltis et al. (1983).

| Enzyme | E.C. # | Results |  |  |  |
| --- | --- | --- | --- | --- | --- |
| PGI | 5.3.1.9 | POLYMORPHIC, BUT NOT SCORABLE |  |  |  |
| PGM | 2.7.5.1 | POLYMORPHIC, BUT NOT SCORABLE |  |  |  |
| SKdh | 1.1.1.25 | MONOMORPHIC |  |  |  |
| ALD | 4.1.2.13 | MONOMORPHIC |  |  |  |
| SOD | 1.15.1.1 | MONOMORPHIC |  |  |  |
| IDH | 1.1.1.42 | MONOMORPHIC |  |  |  |
| PRO | PROTEIN | MONOMORPHIC, MULTIPLE BANDS |  |  |  |
| ENZYMES TESTED, BUT NOT ADEQUATELY RESOLVED |  |  |  |  |  |
| ENZYME | E.C. # | ENZYME | E.C. # | ENZYME | E.C. # |
| CAT | 1.11.1.6 | FK | 2.7.1.4 | DIA | 1.6.2.2 |
| PPO | 1.10.3.2 | 6PG | 1.1.1.44 | ACON | 4.2.1.3 |
| GalDH | 1.1.1.48 | MDH | 1.1.1.37 | ALD | 4.1.2.13 |
| PEP | 3.4.15.1 | G6PDH | 1.1.1.49 | FUM | 4.2.1.2 |
| AAT | 2.6.1.1 | AKP | 3.1.3.1 | FDP | 3.1.3.11 |
| HK | 2.7.1.1 | ACP | 3.1.3.2 | PFK | 2.7.1.11 |
| ME | 1.1.1.40 | EST | 3.1.1.1 | MANDH | 1.1.2.2 |
| PER | 1.11.1.7 | SDH | 1.3.99.1 | MuEST | 3.1.1.1 |
| ADK | 2.7.4.3 | SucDH | 1.3.99.1 | B-GAL | 3.2.1.23 |
| LAP | 3.4.11.1 | LDH | 1.1.1.27 | AMY | 3.2.1.1 |
| TPI | 5.3.1.1 | ADH | 1.1.1.1 | GlutDH | 1.4.1.2 |

## Discussion

Different strains of *Pilayella littoralis* have adapted to the wide range of seawater temperatures found throughout the geographic distribution of the species. Within Nahant Bay, Massachusetts, growth responses to various temperature regimes in culture also differed between the unique free-living and attached forms. In addition, the Nahant Bay free-living form has diverged in morphology and reproductive characteristics from the attached form.

Ecotypic temperature differentiation was demonstrated for the Greenland isolate of *Pilayella littoralis.* Compared to Massachusetts and Connecticut, the Greenland isolate is narrowly adapted to temperature, and exhibits maximum growth at colder temperatures (Figure 2). Interestingly, maximum growth in culture occurred at temperatures above that ever reached by seawater in the region. The intertidal zone of Disko Island, Greenland, is free of ice only in the summer months (June to October), and the growth of *P. littoralis* correlates with seawater and intertidal temperatures during this period. Air temperatures, in particular, reach or exceed 10° C during the summer, and the growth of the alga may reflect ambient air temperature rather than seawater temperature. The dark basaltic substratum of these isolates would also heat up in the sun at low tide in summer months (intertidal range 0.3 - 2.8 m). Examples of temperature optima for growth that are considerably above ambient seawater conditions are found in *Laminaria* spp. (Luning, 1980), a sublittoral alga, and *Ectocarpus siliculosus* (Dillw.) Lyngb. (Bolton, 1983), an intertidal and sublittoral species.

Further evidence of temperature ecotypic variation in *Pilayella littoralis* is found in the narrow growth response of the Nahant Dread Ledge isolate. The Dread Ledge population is perennial (pers. comm. R.T. Wilce) and growth is poor at 20° C (Figure 3). Since the population reproduces by motile cells, selection may act on these propagules to fine-tune growth responses to immediate shifts in ambient temperature. Also, the maximum summer water temperature is significantly below 20° C (Figure 2). Diatoms, another form of free-living algae, show genetic changes that occur within populations on a seasonal basis (Gallagher, 1980). With typically large numbers in diatom populations, and rapid turnover rates of individuals, selection occurs to change the genetic composition of the population as it moves through the seasons. Epilithic isolates of Dread Ledge *P. littoralis* may exhibit a similar phenomenon, where genetic variation within the population rather than phenotypic plasticity accounts for its persistence year-round.

The growth response curve of epiphytic *Pilayella littoralis* from Connecticut was similar to that of *P. littoralis* from Nahant Bay. The Connecticut isolate, however, exhibited a shift in reproductive response that was unique to attached forms of the species. Reproduction by motile spores was limited almost entirely to 20° C. The Connecticut population persists throughout the summer and is most abundant when water temperatures are warmest (pers. comm. J. Foertch). Whether reproduction in the field occurs at the same temperature as in culture is unknown. The Connecticut isolate was held in culture for one (long-day growth study) to three months (short-day study) before use as experimental material. Additional growth studies are needed to confirm that this reproductive response is genetic.

Seaweeds throughout their geographical range do not always exhibit ecotypic temperature variation. For example, Bolton and Luning (1982) tested eight populations of *Laminaria* spp. for evidence of ecotypic adaptation and found no significant interpopulation growth differences in culture, despite large differences in temperature between collecting sites. Rietema and van den Hoek (1984) failed to find latitudinal adaptations to temperature in *Dumontia contorta* (Gremlin) Ruprecht, collected through much, but not all, of the alga’s northeastern Atlantic latitudinal range. Similarly, McLachlan and Bird (1984) did not find temperature ecotypes in species of *Gracilaria*.

The answer to how ecotypic differentiation arose in free-living *Pilayella littoralis* remains speculative. At some point in the past, the free-living population of *P. littoralis* in Nahant Bay originated from propagules of the attached form (Wilce et al. 1982). However, when and how propagules from attached forms of the species gave rise to the free-living population is unknown, though the nuisance population has been in Nahant Bay for at least 120 years (Wilce et al., 1982). If individuals from attached populations do not currently contribute propagules to the free-living population, then by clonal reproduction the free-living population would consist of individuals all sharing the same genotype, or only a few genotypes might exist that represent a hypothetical founder population. Alternatively, the free-living population might be highly polymorphic, with genetic variation a result of continual input from attached populations. Additionally, somatic mutations within the free-living population might accumulate over time.

If considerable genetic variability exists in attached populations of *Pilayella littoralis* (a reasonable assumption), then it’s possible that some individuals are preadapted to the free-living habit and ultimately contribute to maintaining the free-living population. Broad growth response to temperature may be one factor that predisposes typically ephemeral attached *P. littoralis* to survive as a perennial free-living form. On the other hand, attached *P. littoralis* exhibits determinate growth. The species produces by motile reproductive cells following the differentiation of vegetative cells into reproductive cells. The release of motile cells from a unilocular or plurilocular reproductive organ is the end of the cell line that leads up to their production; vegetative cells reproduce and die. Free-living *P. littoralis* lacks reproduction by motile cells. Consequently, growth in this form of the species is indeterminate and large clones may be extensive in the population. Whether indeterminate growth is characteristic only of the free-living habit or also present as part of the genetic background of attached *P. littoralis* is unknown.

Culture studies suggest that propagules from attached *Pilayella littoralis* are not transformable to the free-living form of the species (Figure 15). Bubbled cultures that, in part, mimicked the free-living environment did not transform attached plants into the free-living ball form even after eight months in culture. Similarly, under static conditions, free-living *P. littoralis* could not be converted into algae with the morphology and reproductive characteristics of the attached form. It is possible that the culture conditions were not sufficient to induce the transformation of these two forms. Or, despite the prolonged duration of this experiment, the culture studies were terminated before the algae had a chance to develop transformed characteristics.

However, if transformation from attached to free-living morphology is a lengthy process, and if attached algae frequently contribute individuals to the free-living population, then recognizable attached forms should be evident among the free-living community. Attached forms of the species are easily distinguished by their abundant reproductive organs, extensive cabled central axis, and basal-distal polarity. Attached forms are rarely found among free-living P. *littoralis*, but when discovered, they exhibit characteristics of recently detached material. Notably, some isolates of free-living material retain the capacity to produce reproductive organs. Whether the ability to produce unilocular or plurilocular organs is never wholly lost or whether these isolates represent a transitional stage in an adaptive process leading to the free-living form of the species is unknown.

Isozyme data could resolve questions concerning the genetic structure of free-living and attached populations in Nahant Bay, Massachusetts. However, based on the preliminary data of this study, the free-living and attached populations of *Pilayella littoralis* in Nahant Bay are not distinguishable at the enzyme level. One variable enzyme system (PGI) of two alleles at one locus, with little polymorphism, is insufficient to determine the genetic relationship between the two forms of *P. littoralis.* The most that can be said is that since some variability was observed in the free-living population, it can be ruled out that this form of the alga represents one large genetically identical clone. Results of the common garden experiments reported above suggest that genetic differentiation has occurred between attached and free-living *P. littoralis.* The electrophoretic data are not incompatible with the culture results. The conclusion of no detectable difference at the enzyme level between Nahant Bay populations of these algae is based on a lack of data to reject the null hypothesis.

The ability to extract enzymes that can be resolved on starch or acrylamide gels has proven to be a significant obstacle in studying seaweed population genetics (Cheney, 1985). This research did not prove an exception. Few seaweeds have been examined electrophoretically (for reviews, see Innes, 1984; Chapman, 1986). For example, in cultured populations of *Porphyra yezoensis* (Miura et al., 1979), polymorphism, as percent loci polymorphic, was 0%. In *Enteroaorpha linza,* polymorphism was 100% (Innes, 1984). The only other studies that have included allele frequencies are for *Codium fragile* (4 of 12 loci polymorphic; Malinowski, 1974), *Euchuema* sp. (4 of 7 loci polymorphic; Cheney and Babbel, 1978), and *Porphyra yezoensis* (wild populations with 6 of 8 loci polymorphic; Miura et al., 1979). Considerably more data are needed before generalizations can be made regarding genetic variability in *P. littoralis* or seaweeds in general.

Unattached seaweed populations with attached counterparts are not uncommon (reviewed by Norton and Mathieson, 1983). Little is known, however, regarding the development of unattached seaweed populations, especially about environmentally or genetically induced differences that occur following detachment from native substrata. A characteristic of many unattached algae that may have significant genetic implications is the absence of reproductive organs (Moore, 1950, King, 1981; re. *Hormosira banksii* (Turner); Gibb, 1957; South and Hill, 1970; re. *Ascophyllum nodosum;* Gorden et al., 1985; re. *Cladophora montagneana* KOtz.; Wilce et al., 1982; re. *Pilayella littoralis).* For many taxa, except for periodic “seeding” from attached populations, vegetative fragmentation is the primary way unattached seaweed populations reproduce and give rise to new individuals. Consequently, unattached seaweed populations may be reproductively isolated from their attached progenitors (no gene flow) and are therefore free to respond to the unique selection pressures of the free-living environment.

Once a free-living population of seaweed becomes established as a perennial population, somatic mutations may provide the genetic variation upon which selection acts to isolate this form of the species. Growth rates in the free-living *Pilayella littoralis* population are high, and turnover times for individual free-living balls are rapid. These traits all favor the spread of favorable mutations through the population. Free-living *P. littoralis* may be in the nascent stages of speciation.

If phenotypic plasticity is sufficient to respond to environmental stresses associated with transfer from an attached to the free-living habit, then free-living populations can be maintained by input from nearby attached populations. The term ecad has been used to describe morphological variation seen in many unattached algal populations (Gibb, 1957; Brinkhuis, 1976; Russell, 1986), where selection acts on the character of phenotypic plasticity *(sensu* Bradshaw, 1965) to produce morphologies distinctive from the typical form of the species. For example, *Ascophyllum nodosum* ecads develop from fragments of individuals that are transported, usually, to saltmarsh environments (Gibb, 1957; Brinkhuis, 1976). The development of morphologies different from typically attached individuals depends on the interaction between abiotic and biotic factors. In particular, the apical cell of unattached *A. nodosum* reverts to juvenile form (three-sided from four-sided), which has been suggested as responsible for the distinctive character of various ecads of the species (Fritsch, 1945; Baker, 1950; Moss, 1956). *A. nodosum* is a particularly well-studied seaweed; comparable information on other unattached seaweed populations is unavailable.

Unlike *Ascophyllum nodosum, Pilayella littoralis* has simple thallus construction, diffusive growth, and rapid turnover of individuals in the population. This suggests that interactive effects between the environment and P. *littoralis* do not result in modified developmental processes as in *A. nodosum.* This idea is supported by the inability to transform the species in laboratory experiments and the stability of the free-living form under a wide range of temperatures, irradiances, and photoperiod conditions.

When considering ecotypic variation among seaweeds it is important to consider multiple traits (Russell, 1986). For example, the branching pattern in *Ectocarpus siliculosus* does not display ecotypic variation (Russell, 1967), but copper tolerance does (Russell and Morris, 1970). Similarly, the blade morphology of *Macrocystis pyrifera* (L.) C. Agardh is plastic, but the rate of blade initiation is not (Druehl and Kemp, 1982). Further, more than one set of experimental manipulations may be required to confirm that differences between algal types are ecotypic. For example, for attached and free-living *Pilayella littoralis,* both transplant and growth studies in the laboratory provide more convincing evidence of ecotypic differentiation than either experiment offers alone.

As shown by a series of common garden growth experiments, these studies have determined that genetic differentiation exists between Nahant Bay free-living and attached *Pilayella littoralis,* and among geographic isolates of the species. The extent of genetic differentiation between the two Nahant Bay forms is unknown. However, suppose that specific genotypes within attached populations are preadapted to survive in the free-living state; then, the free-living population may simply represent a subset of overall genetic variability within the attached form of the species. Once an individual establishes free-living existence, it is effectively immortal. Indeterminate growth and clonal reproduction by fragmentation favor the accumulation of neutral or favorable somatic mutations. Some individuals in the free-living population may, therefore, be genetically distinct from the gene pool of its origin. Therefore, genetic markers that are not dependent on phenotypic expression are needed to learn more about the genetics of P. *littoralis* in Nahant Bay. For example, isozymes or restriction fragment length polymorphisms may provide the resolution necessary to answer whether the free-living population is one large clone, many clones, or if the attached population significantly contributes to maintaining the free-living form in Nahant Bay.

## Acknowledgments

I (SLM) thank my Ph.D. advisor Robert T. Wilce for his support and friendship. After RTW recently passed, I discovered the absence of additional ecological work on free-living *Pilayella littoralis* since his seminal studies on the alga and completing my thesis. Therefore, to honor RTW, I hope readers find the work interesting and relevant. I also dedicate the research to Robert Vadas, who taught me experimental ecology early in my graduate career. I carry those lessons with me today. I thank Charlie Yarish who shared culture techniques. Andrew Davis and Timothy Briggs helped with sample collections and diving. RTW received a grant from the Metropolitan District Commission: Division of Parks, Engineering, and Construction to support this work. SLM received funding from the Woods Hole Trust Fund, University of Massachusetts.

